# CHK1-CDC25A-CDK1 regulate cell cycle progression in early mouse embryos to protect genome integrity

**DOI:** 10.1101/2022.11.13.516318

**Authors:** Knoblochova Lucie, Duricek Tomas, Vaskovicova Michaela, Zorzompokou Chrysoula, Rayova Diana, Ferencova Ivana, Baran Vladimir, Richard M. Schultz, Eva R. Hoffmann, Drutovic David

**Affiliations:** Institute of Animal Physiology and Genetics of the Czech Academy of Sciences, 277 21 Libechov, Czech Republic; Faculty of Science, Charles University,128 00 Prague, Czech Republic; Institute of Animal Physiology, Centre of Biosciences, Slovak Academy of Sciences, 040 01 Kosice, Slovakia; Department of Biology, University of Pennsylvania, Philadelphia, PA 19104, USA; Department of Anatomy, Physiology, and Cell Biology, School of Veterinary Medicine, University of California, Davis, CA 95616, USA; DNRF Center for Chromosome Stability, Department of Cellular and Molecular Medicine, Faculty of Health and Medical Sciences, University of Copenhagen, 2200 N Copenhagen, Denmark

**Keywords:** CDC25A phosphatase, CDK1 kinase, cell cycle regulation, CHK1 kinase, early mouse embryos

## Abstract

After fertilization, remodeling of the oocyte and sperm genomes is essential to convert these highly differentiated non-dividing transcriptionally quiescent cells into early cleavage-stage transcriptionally active totipotent blastomeres. This developmental transition is accompanied by cell cycle adaptation such as lengthening or shortening of the gap phases G1 and G2. However, regulation of these cell cycle changes is poorly understood, especially in mammals. Checkpoint kinase 1 (CHK1) is a protein kinase that regulates cell cycle progression in somatic cells. Here, we show that CHK1 regulates cell cycle progression in early mouse embryos by restraining CDK1 kinase activity due to CDC25A phosphatase degradation. CHK1 kinase also ensures the long G2 phase needed for genome activation and reprogramming gene expression in 2-cell stage mouse embryos. Last, *Chk1* depletion leads to DNA damage and chromosome segregation errors that result in aneuploidy and infertility.

## INTRODUCTION

Preimplantation development is associated with several critical developmental changes that include initial embryonic cell divisions, zygotic genome activation, compaction, and blastocyst formation. The egg and sperm are highly differentiated cells whose chromatin structure has dramatically different epigenetic landscapes (Eckersley-Maslin et al, 2018; Kobayashi et al, 2012; Sankar et al, 2020). Following fertilization these differences are resolved largely during the first cell cycle as the paternal and maternal genomes are remodelled to provide a chromatin structure that supports the dramatic reprogramming of gene expression associated with genome activation (Eckersley-Maslin et al, 2018). These processes are linked to significant cell cycle changes and regulated by maternally-derived proteins and mRNAs stored in the egg until transcription from the embryonic genome is established.

The first two embryonic cell divisions are about 18-20 h long in mouse and human embryos. Duration of individual cell cycle phases in initial embryonic cell divisions differs mainly in the length of the gap phases. In contrast to the first mitotic G1 phase, which lasts about 5-12 h, dependent on the genetic background of mice, 2-cell stage embryos spend only 1-2 h in the G1 phase during the second cell division. Lengthening the first G1 phase likely provides sufficient time to form both pronuclei, which requires chromatin decondensation of the egg and sperm nuclei and chromatin remodelling (Artus & Cohen-Tannoudji 2008; Palmer & Kaldis 2016; Radonova et al, 2019). In mice, transcription of the embryonic genome starts during the first S phase in the zygote and slowly increases from G2 (“minor” genome activation) to a peak in G2 in late 2-cell stage embryos (“major” genome activation) (Fugaku Aoki et al, 1997; Braude 1979; Flach et al, 1982; Molls et al, 1983). The second division is characterized by the almost virtual absence of a G1 phase and an dramatic lengthening of G2/M transition (approximately 12-16 h) compared to 1-8 h for G2/M in the first division (Abramczuk & Sawicki 1975; Domon 1980; Howlett & Bolton 1985; Krishna & Generoso 1977; Molls et al, 1983; Palmerola et al, 2022). This extended G2 phase likely provides sufficient time for expression of products of major genome activation to accumulate to sufficient concentrations and thereby support continued development (Flach et al, 1982; Molls et al, 1983). Genome activation in other mammals is delayed to the 4-16-cell stage, although there is evidence for earlier transcription (Xue et al, 2013). Mechanisms underlying regulation of cell cycle duration in the initial divisions, in particular the transition from a long G1 and short G2 in zygotes to a short G1 long G2 in 2-cell embryos, remain largely unknown.

Cell cycle progression is largely regulated by cyclin-dependent kinases (CDK). CDKs are inhibited by WEE/MYT kinase-mediated phosphorylation of T14 or Y15. To activate CDKs, these inhibitory phosphorylations are removed by CDC25 phosphatases (Enoch & Nurse 1990). Although CDKs in mammals have many paralogues - *Cdk1, Cdk2, Cdk4*, and *Cdk6*, CDK1 is the only essential one (Santamaría et al, 2007). CDK1 can compensate for the other CDKs and is indispensable for meiotic (Adhikari et al, 2012) and mitotic progression (Santamaría et al, 2007). *Cdc25* phosphatase in mammals has three paralogues, *Cdc25a, Cdc25b*, and *Cdc25c* (Galaktionov & Beach 1991; Nagata et al, 1991; Sadhu et al, 1990), with CDC25A being the only essential one (Chen et al, 2001; Ferguson et al, 2005; Lee et al, 2009; Lincoln et al, 2002; Ray et al, 2007).

Genome integrity relies on communication between the cell cycle progression, checkpoint activation, and DNA repair pathways. DNA damage activates the DNA damage response (DDR) and cell cycle checkpoint signaling, which retards cell cycle progression to ensure sufficient time to repair DNA damage. Interestingly, the ability of cell cycle checkpoint signaling proteins to modulate cell cycle duration is utilized for early development (Artus & Cohen-Tannoudji 2008). However, little is known about DDR signaling and checkpoint activation in early embryos. It should be noted that incompletely replicated DNA caused by replication stress during human embryonic development is a major cause of an increased incidence of aneuploidy in human embryos (Palmerola et al, 2022).

ATR and CHK1 are DDR kinases that regulate cell cycle progression by maintaining replication checkpoint signaling (Eykelenboom et al, 2013; Kalogeropoulos et al, 2004; Lebrec et al, 2022; Saldivar et al, 2018; Zachos et al, 2005). The replication checkpoint prevents mitosis before DNA replication is finished (Enoch & Nurse 1990; Hartwell & Weinert 1989). CHK1, not ATR, is the DDR limiting kinase in the zygote (Ladstätter & Tachibana-Konwalski 2016). In addition, CHK1 depletion or inhibition results in premature mitosis and chromosome fragmentation in mice (Fishler et al, 2010; Lam et al, 2004; Niida et al, 2005; Peddibhotla et al, 2009), avian (Zachos et al, 2007, 2003, 2005) and human (Branigan et al, 2021; Durkin et al, 2006; Niida et al, 2005; Syljuåsen et al, 2005) somatic cells. CHK1 also regulates cell cycle progression during embryonic development in zebrafish (Dalle Nogare et al, 2009; Zhang et al, 2014), *Drosophila* (Deneke et al, 2016) and *Xenopus* (Petrus et al, 2004; Shimuta et al, 2002). Moreover, CHK1 inhibition or depletion in human (Palmerola et al, 2022) and mouse embryos (Ju et al, 2020; Liu et al, 2000; Muralidharan et al, 2020; Takai et al, 2000) exacerbates the incidence of aneuploidy and induces genome fragmentation.

Although the mechanism underlying the role of CHK1 in cell cycle checkpoint signaling is well understood in somatic cells, little is known regarding the function of CHK1 signaling in regulating embryonic cell division and genome integrity in early mammalian embryos. Here, we show that maternal *Chk1* depletion in mice causes acceleration of cell cycle progression accompanied by severe chromosome segregation errors, especially in 2-cell embryos. Live-cell imaging revealed that maternal *Chk1* depletion shortens all cell cycle phases in 2-cell embryos, most notably, the long G2 phase needed for genome activation. The shortening is due to premature CDK1 kinase activation caused by stabilization of CDC25A phosphatase.

## RESULTS

### Maternally-expressed *Chk1* regulates duration of the first and second cleavage divisions

We previously reported that maternally-expressed *Chk1* is not required for oocyte maturation but is essential for preimplantation development in mouse (Ruth et al, 2021); maternally-depleted *Chk1* embryos show impaired development to the blastocyst stage, massive genome fragmentation at the 3-4- or 5-8-cell stage, and a lack of liveborn pups. To further understand the maternal contribution of CHK1 during preimplantation development, we used the same mouse model where Cre recombinase is expressed using the zona pellucida-3 (*Zp3)* promoter to remove floxed exon 2 of *Chk1* in growing oocytes (Lam et al, 2004; Ruth et al, 2021) (**Supplemental Fig. S1A, B**). Females lacking *Chk1* in oocytes were mated with wild-type (WT) males to produce maternally-depleted *Chk1* embryos (*Chk1 mKO*) (**Supplemental Fig. S1C**). Controls were litter mates carrying a floxed exon 2 but not Cre recombinase, which results in CHK1 expression as in WT (*Chk1 WT*) (**Supplemental Fig. S1C**). Immunoblot analysis showed that CHK1 protein was not detected in WT sperm (**Supplemental Fig. S1D**), thereby minimizing any paternal contribution to *Chk1* mKO embryos. We observed low expression of full-length and truncated CHK1 in *Chk1 mKO* zygotes (**Supplemental Fig. S1E**). The truncated CHK1 version may have arisen by initiating translation from the first methionine reside encoded in exon 3, which would yield CHK1 lacking the initial 41 amino acids (*Chk1 Δ*^*2*^) and giving rise to a species with a predicted band size of 50 kDa, which was observed on the immunoblot (**Supplemental Fig. S1B, E**).

To analyze the role of CHK1 in cell cycle progression during preimplantation development, bright-field images of live, *Chk1 WT* and *Chk1 mKO* embryos were collected every 30 min and individual embryos analyzed for cell cycle progression (**Fig. 1A, B**) (Artus & Cohen-Tannoudji 2008; Nothias et al, 1995). At 114 h after hCG administration, 53% of *Chk1 WT* zygotes formed blastocysts, compared to 12% of *Chk1 mKO* zygotes (**Fig. 1C**). In addition, *Chk1 mKO* embryos exhibited an increase in cellular fragmentation (**Fig. 1B, C**). Blastocysts that formed from *Chk1 mKO* embryos could develop further because some dead pups, but no live pups, were born (Ruth et al, 2021). We confirmed these results by culturing zygotes in the presence of 250 nM CHIR-124, a CHK1 inhibitor (Ju et al, 2020), and observed embryonic arrest and failure to form blastocysts (**Supplemental Fig. S1F**). This phenotype was observed both in C57J/BL6 and CD-1 mouse strain backgrounds (**Supplemental Fig. S1F**).

**Figure 1.**
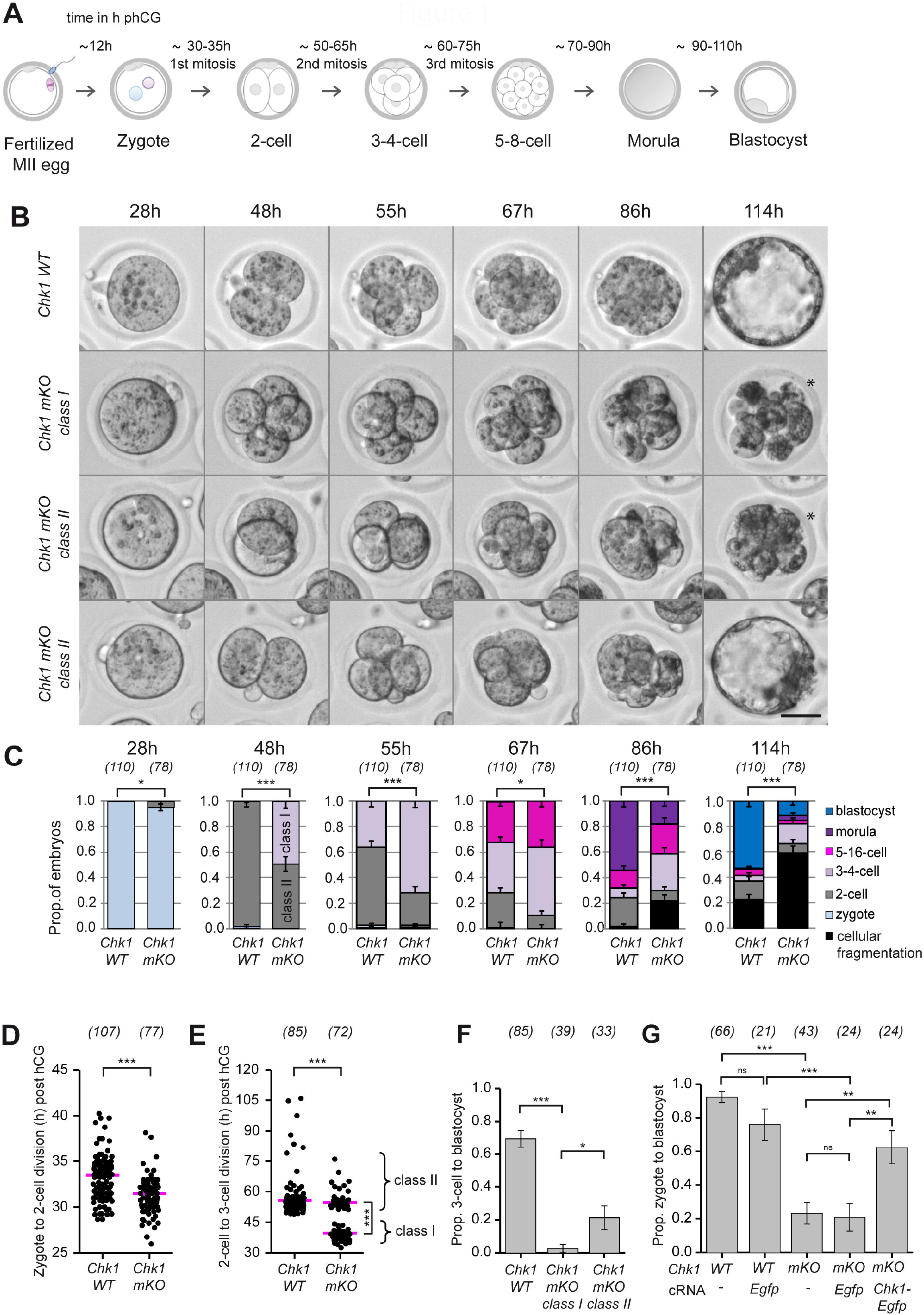
*Chk1 mKO* embryos display accelerated first and second embryonic division and compromised development. **A** Schematic of embryonic development using bright-field time-lapse imaging. Time (h) is relative to h post hCG administration. **B** Representative still images from bright-field time-lapse imaging of preimplantation development of *Chk1 WT, Chk1 mKO class I*, and *Chk1 mKO class II* embryos from 28 to 114 h post hCG administration. Strain background: C57J/BL6, with small part CD-1. An asterisk marks fragmented embryos. Scale bar 30 μm. **C** Data from bright-field time-lapse imaging (**B**) was used to quantify the proportion of *Chk1 WT*, and *Chk1 mKO* embryos at different developmental stages. At 48 h, the proportion of *Chk1 mKO class I*, and *Chk1 mKO class II* embryos is shown. The numbers in parentheses show the number of embryos imaged for each genotype. The data were pooled from three independent experiments. *p<0.05, and ***p<0.001 (Cochran-Armitage Trend Test, any trend). **D, E** Data from bright-field time-lapse imaging (**B**) was used for quantification the division time (h) from zygote to 2-cell (**D**) and from 2-cell to 3-cell (**E**) in *Chk1 WT*, and *Chk1 mKO*, embryos. Distribution of *Chk1 mKO class I*, and *Chk1 mKO class II* is shown in **E**. The median is shown. ***p<0.001 relative to the *Chk1 WT* group (Mann-Whitney U test, two-sided), ***p<0.001 relative to the *Chk1 mKO class I* group (Kolmogorov-Smirnov Test, 2-sided). **F** Data from bright-field time-lapse imaging (**B**) was used to quantify the proportion of 3-cell embryos that developed to the blastocyst stage. Error bars denote standard error of proportion. *p<0.05, and ***p<0.001 (Fisher’s Exact Test). **G** Proportion of *in vitro* blastocyst development in wild type (WT), and *Chk1 mKO* (mKO) zygotes microinjected with *Chk1*-*Egfp* or *Egfp* cRNA. Error bars denote the standard error of proportion. The data were pooled from two experiments, one with *Chk1 mKO* and *Chk1*-*Egfp* cRNA microinjection and the other with *Chk1 mKO* and *Egfp* cRNA microinjection. In both experiments, the embryos from one group were divided into microinjected and non-microinjected (serving as controls) arms. n.s. not significant, **p<0.01, and ***p<0.001 (Fisher’s Exact Test).

Bright-field time-lapse data revealed that *Chk1 mKO* zygotes divided to the 2-cell stage about 2 h earlier than *Chk1 WT* and these mutant embryos also divided sooner (measured as 2-cell to 3-cell stage division) (**Fig. 1D, E**), with 54% dividing 13 h earlier than *Chk1 WT* (hereafter referred to as *Chk1 mKO class I*) (**Fig. 1C, E**). The 46% of *Chk1 mKO* embryos that divided with a similar timing as *Chk1 WT* are referred to as *Chk1 mKO class II* (**Fig. 1C, E**). *Chk1 mKO class I* showed both acceleration in the first (zygote to 2-cell) and second (2- to 3-cell) divisions that was accompanied by a near absence of blastocyst development. Although, *Chk1 mKO class II* embryos displayed the *Chk1 WT* like timing of divisions, only about 20% developed to form blastocysts (**Fig. 1F, Supplemental Fig. S1G**). In addition, a positive correlation was observed between the timing of the first and second embryonic division in *Chk1 mKO class I* embryos (**Supplemental Fig. S1H**). These changes in timing of cell division were due to loss of maternal CHK1 because microinjecting *Chk1 mKO* zygotes with *Chk1-Egfp* cRNA, but not *Egfp* cRNA, rescued the mutant phenotype (**Fig. 1G**).

To ascertain whether differences in the residual amount of CHK1 in *Chk1 mKO* embryos was the basis for the *class I* or *class II* phenotype, we performed an CHK1 immunoblot analysis using embryos harvested 47-48 h post-hCG (**Supplemental Fig. S1I**). *Chk1 mKO* embryos showed a slightly higher amount of CHK1 in *Chk1 mKO class II* embryos compared to *Chk1 mKO class I* embryos, suggesting that the *Chk1 mKO class II* phenotype was caused by some residual CHK1 protein compared to *Chk1 mKO class I*. Because we did not detect expression of the truncated CHK1 version in these two classes in contrast to *Chk1 mKO* zygotes (**Supplemental Fig. S1B, E**), it was unlikely that the truncated form plays a significant role in *Chk1 mKO* embryos. To further investigate the distinction between *Chk1 mKO class I and class II* embryos, we assessed the proportion of *Chk1 mKO class I* embryos in individual mice. Most mice showed 50-80% of *Chk1 mKO class I* embryos (**Supplemental Fig. S1J**). We conclude that full-length maternally-derived CHK1 regulates the timing of initial embryonic divisions and is required for preimplantation development to blastocyst.

### CHK1 kinase does not participate in replication checkpoint

In somatic cells, CHK1 kinase prevents mitotic entry until DNA replication is completed (Kalogeropoulos et al, 2004; Lebrec et al, 2022; Saldivar et al, 2018; Zachos et al, 2005). CHK1 kinase also regulates S phase progression, including replication origin firing, replication fork speed progression, and DNA damage response in S phase (Dandoulaki et al, 2018; Michelena et al, 2019; Niida et al, 2005; Syljuåsen et al, 2005). To determine whether the accelerated zygote to 2-cell division occurs prior to completion of DNA replication, we assayed DNA replication in *Chk1 mKO* embryos by DAPI staining. Because CHK1 depletion increases DNA damage (Dandoulaki et al, 2018, Niida et al, 2005), we also assessed in the same embryo formation of DNA double-stranded breaks (DSBs) by immunofluorescence detection of phosphorylated histone H2AX on S139 (γH2AX) (**Supplemental Fig. S2A**) (Mah et al, 2010; Sedelnikova et al, 2002). DNA replication triggers transient DDR signaling with a high γH2AX signal in S phase (Xu et al, 2015). Accordingly, embryos were treated with a 60 min pulse of the thymidine analog 5-ethynyl-2-deoxyuridine (EdU) to exclude zygotes in S phase from analysis (**Supplemental Fig. S2A**) and fixed zygotes when most *Chk1 WT* embryos were in G2 (**Fig. 1A**) (Artus & Cohen-Tannoudji 2008), i.e., at 28 h post-hCG administration. EdU staining also allowed us to monitor cell cycle progression.

We did not observe massive DNA underreplication in *Chk1 mKO* zygotes compared to *Chk1 WT* zygotes (**Supplemental Fig. S2B**). Analysis of zygotes in G2 revealed a similar and relatively broad distribution of γH2AX foci in *Chk1 WT* and *Chk1 mKO* (**Supplemental Fig. S2C**). As expected from bright-field imaging (**Fig. 1D**), we observed a significantly different distribution of cell cycle phases in *Chk1 mKO* zygotes compared to *Chk1 WT* (**Supplemental Fig. S2D**). The significant decrease in the percentage of S-phase *Chk1 mKO* zygotes suggests a shortened G1 and/or S phase in zygotes lacking maternal CHK1. Taken together, *Chk1 mKO* zygotes show neither massive DNA underreplication nor elevated DSBs but do exhibit altered cell cycle progression and accelerated G2/M entry.

To gain insight for the basis for the accelerated second division in *Chk1 mKO* embryos, we examined DNA content, DSBs formation, and cell cycle progression in late 2-cell stage embryos. We fixed the embryos at 50 h after hCG administration, when *Chk1 WT* and *Chk1 mKO class II* embryos should be in the 2-cell stage, but *Chk1 mKO class I* should be in the 3- or 4-cell stage (**Fig. 1B, C**). Consistent with our previous results, *Chk1 mKO* showed *class I* (3-4-cell stage) and *class II* (2-cell stage embryos) (**Supplemental Fig. S2E, F**) distributions, with *Chk1 mKO class I* embryos showing significant increases in genome fragmentation and micronuclei formation (**Supplemental Fig. S2E, G**). A 60-minute EdU pulse revealed that all 2-cell embryos were in G2 (**Supplemental Fig. S2E**). In addition, DNA content (assayed by DAPI staining) in 2-cell stage *Chk1 mKO class II* embryos was similar to 2-cell stage *Chk1 WT* (**Supplemental Fig. S2H**), whereas DSBs formation (assayed by γH2AX detection) was greater than *Chk1 WT* (**Supplemental Fig. S2I**). Thus, *Chk1 mKO class II* embryos do not show massive DNA underreplication but exhibit increased DNA damage. Our results suggest that CHK1 does not participate in the replication checkpoint in early cleavage-stage mouse embryos because DNA content in *Chk1 mKO* was similar to *Chk1 WT* (**Supplemental Fig. S2B, H**).

Apart from CHK1, WEE/MYT protein kinases also participate in replication checkpoints in somatic cells (Enoch & Nurse 1990). To ascertain whether WEE/MYT are involved in the replication checkpoint in early embryos, we used embryos from transgenic *H2b-Egfp* mice to label chromosomes for live imaging (Hadjantonakis & Papaioannou 2004; Mayer et al, 2016) and treated them with the WEE/MYT kinase inhibitor PD0166285 21 h after hCG administration (**Supplemental Fig. S3A**), when the embryos were in late G1 or early S phase (Artus & Cohen-Tannoudji 2008) and then monitored their development by light-sheet microscopy. The treated zygotes entered mitosis, detected by chromosome condensation, as early as 24 h post hCG, when DMSO-treated embryos were in S phase (**Supplemental Fig. S3B, C; Movie M1**). Only 8% of PD0166285-treated embryos completed mitosis and divided to the 2-cell stage (**Supplemental Fig. S3D**). These results show that WEE/MYT kinases regulate mitosis entry from the early or middle S phase, in contrast to CHK1, which does not participate in the replication checkpoint in early embryos.

### CHK1 kinase is required for the long G2 phase in 2-cell stage embryos

We next examined in more detail the accelerated cell cycle progression in 2-cell *Chk1 mKO* embryos. We microinjected *Chk1 WT* and *Chk1 mKO* zygotes with cRNAs for histone *H2b-mCherry* to monitor chromosomes and a modified Fucci(CA) sensor (*mCdt1-Eyfp*) to monitor cell cycle progression (**Fig. 2A**). Because mCDT1-EYFP is degraded during S phase, reappears in G2, and disperses in the cytoplasm after nuclear envelope breakdown (NEBD), we could distinguish the duration of each cell cycle phase in *Chk1 WT* and *Chk1 mKO* 2-cell embryos (**Fig. 2B, C; Movie M2**).

**Figure 2.**
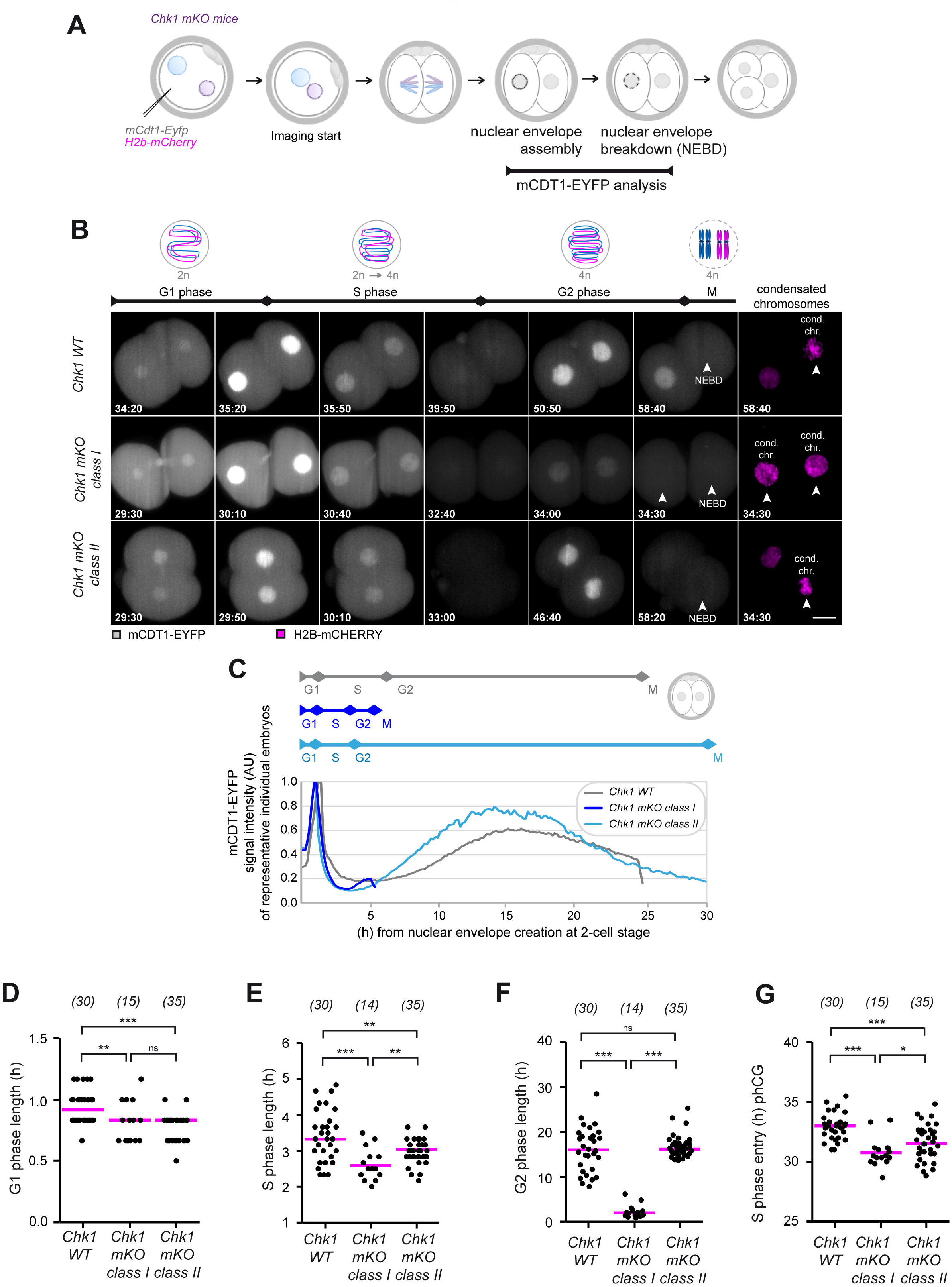
CHK1 kinase regulates cell cycle progression in G1, S, G2, and mitosis. **A** Schematic representation of the experimental design. **B** Representative still images from time-lapse light-sheet imaging of preimplantation development of C*hk1 WT, Chk1 mKO class I*, and *Chk1 mKO class II* embryos. Embryos were microinjected, as shown in **A**. Embryos expressing mCDT1-EYFP (gray) and H2B-mCHERRY (magenta, chromosomes) are shown. Timepoints are relative to time post hCG administration (h). Arrowheads show condensated chromosomes after NEBD. Scalebar 20 μm. Movie M2 related to Fig. 2B: *Chk1 WT* + mCdt1-Egfp Chk1 mKO class I + mCdt1-Egfp Chk1 mKO class II + mCdt1-Egfp **C** Representative line charts of fluorescence signal intensity of mCDT1-EYFP sensor in individual 2-cell stage embryos from time-lapse light-sheet imaging (as shown in **B**) from nuclear envelope formation to nuclear envelope breakdown. Above is a schematic depiction of the cell cycle phase length derived from the line chart. **D, E, F, G** Data from time-lapse light-sheet imaging (**B**) was used to quantify the length (h) of G1 (**D**), S (**E**), and G2 phase (**F**), and S phase beginning (h from hCG stimulation) (**G**) of 2-cell stage embryos. The data were pooled from three independent experiments. The numbers in parentheses show the number of embryos per group. The median is shown. Statitstics is as follows: (**D**) **p<0.01 (Equal-Variance T-Test), *** p<0.001 and n.s. not significant (Aspin-Welch Unequal-Variance T-Test). (**E**) *** p<0.001 and **p<0.01 for *Chk1 mKO class I* (Equal-Variance T-Test). **p<0.01 for *Chk1 WT* (Aspin-Welch Unequal-Variance T-Test). (**F**) *** p<0.001, and n.s. non -significant (Kolmogorov-Smirnov Test). (**G**) *** p<0.001, and * p<0.05 (Equal-Variance T Test).

To validate the information about cell cycle progression from the mCDT1-EYFP sensor, we compared these data to cell cycle progression measured by EdU incorporation. Similar results were obtained with both methods, with S phase ending about 39 to 41 h post hCG administration in WT embryos (**Supplemental Fig. S3E, F, G**). The mCDT1-EYFP signal analysis showed that both *Chk1 mKO class I* and *class II* embryos displayed a slightly but significantly shorter G1 phase due to earlier initiation of S phase compared to *Chk1 WT* (**Fig. 2D**). Also, both *Chk1 mKO class I* and *class II* embryos displayed a significantly shorter S phase, with *Chk1 mKO class I* embryos having a shorter S phase than *Chk1 mKO class II* embryos (**Fig. 2E**). In addition, the majority of *Chk1 mKO class I* embryos did not enter premature mitosis from S phase but from G2 (14 out of 15 embryos). Duration of G2 in *Chk1 mKO class I* embryos was also significantly shortened by about 14 h in comparison to *Chk1 mKO class II* embryos, which showed the same G2 duration as *Chk1 WT* embryos (**Fig. 2F**). In the timing post hCG stimulation, *Chk1 WT* embryos entered S phase from 31h to 36h, whereas *Chk1 mKO* embryos entered S phase from 29 to 35 h (**Fig. 2G**). In toto, these results suggest that CHK1 regulates cell cycle progression of all cell cycle phases but most significant, CHK1 is essential for maintaining the long G2 phase in 2-cell embryos.

### CHK1 kinase regulates long G2 in 2-cell stage embryos by restraining CDK1 activity via CDC25A degradation

In some somatic cell lines, *Chk1* deficiency causes premature entry into mitosis due to precocious CDK1-cyclin B activation (Branigan et al, 2021; González Besteiro et al, 2019; Lemmens et al, 2018; Niida et al, 2005). Because the naturally occurring long G2 phase in 2-cell mouse embryos was shortened in *Chk1 mKO* (**Fig. 2C, F**), we asked whether the shortened G2 in *Chk1 mKO* embryos was caused by premature activation of CDK1. Accordingly, we sought to see if partial CDK1 inhibition could rescue the *Chk1 mKO class I* phenotype by allowing 2-cell embryos to progress to M phase with similar timing to *Chk1 WT*. This approach, namely, partially inhibiting CDK1, was successfully used to inhibit resumption of meiosis, which requires CDK1 activation (Solc et al, 2015) (**Supplemental Fig. S4A**). To determine whether partial CDK1 inhibition was also feasible in mouse embryos, we used different concentrations of RO-3306 (Jang et al, 2014; Spencer et al, 2013; Vassilev et al, 2006). Based on these results, we cultured 2-cell embryos in 2.5 µM RO-3306 and followed cell cycle progression by bright-field microscopy. Treatment with 2.5 µM RO-3306, but not DMSO alone, rescued the onset of mitosis in *Chk1 mKO* embryos (**Fig. 3A, Supplemental Fig. S4B**). These findings, which show that partial inhibition of CDK1 rescues M phase timing in 2-cell stage *Chk1 mKO* embryos, suggest that CHK1 restrains CDK1 activity to regulate cell cycle progression.

**Figure 3.**
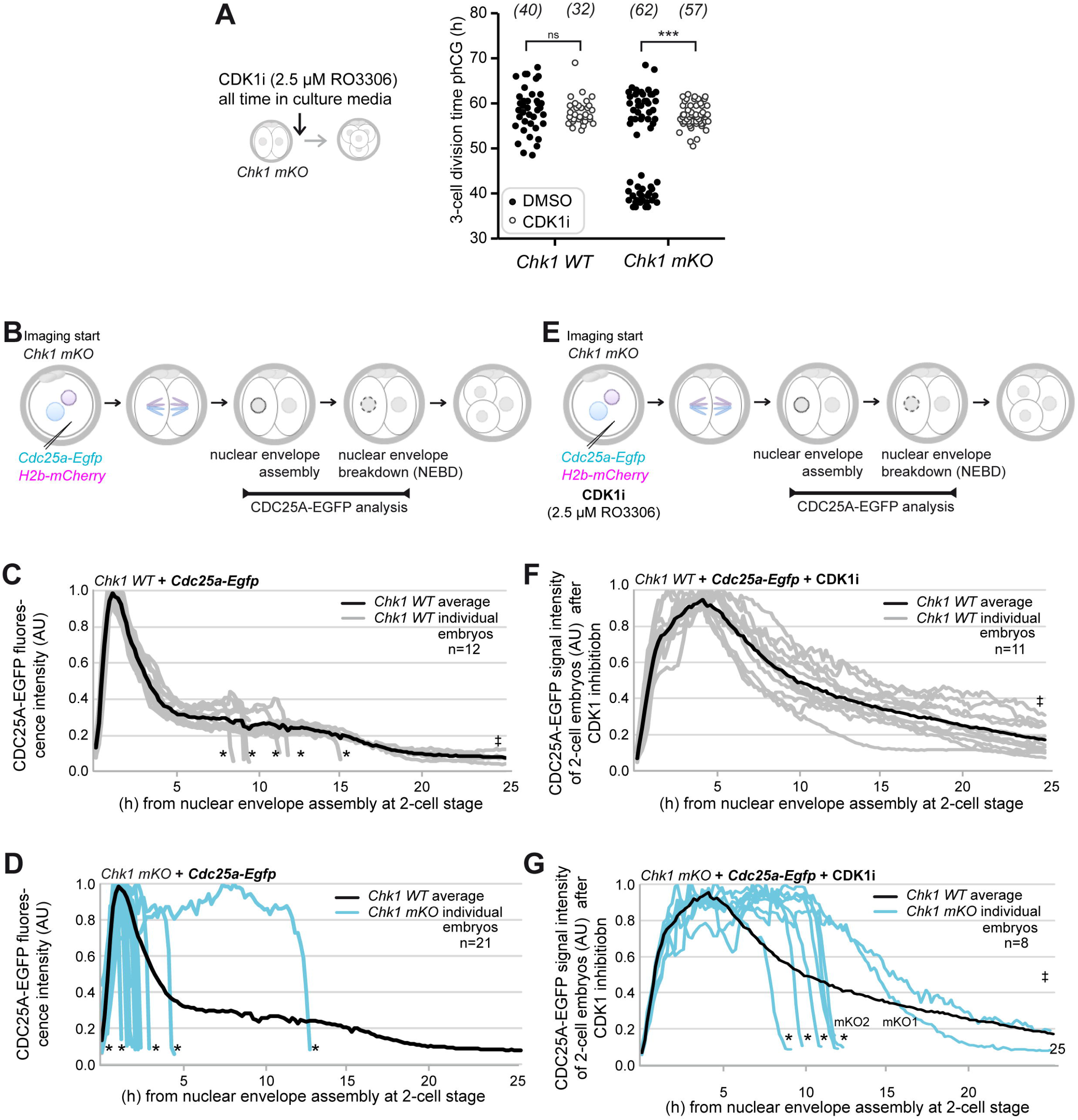
CHK1 kinase regulates long G2 in mouse 2-cell stage embryos by restraining CDK1 activity via CDC25A degradation. **A** Division time (h) from 2-cell to 3-cell stage embryos. *Chk1 WT* and *Chk1 mKO* embryos were treated within 3 h after 2-cell division with 2.5 μM RO3306 (CDK1 kinase inhibitor) or DMSO and imaged by bright-field time-lapse microscopy. The numbers in parentheses denote the number of embryos analyzed per group. Strain background: CD-1 with small part C57J/BL6 and C57J/BL6 with small part CD-1. The data were pooled from two independent experiments. n.s. not significant, and ***p<0.001 (Kolmogorov-Smirnov Test). **B** Schematics of experimental design. **C, D** Line chart of fluorescence signal intensity of CDC25A-EGFP of individual *Chk1 WT* (**C**) and *Chk1 mKO* (**D**) 2-cell stage embryos scanned using light-sheet time-lapse microscopy from nuclear envelope assembly to nuclear envelope breakdown. Embryos were microinjected as shown in **B**. Asterisk mark signal drop after nuclear envelope breakdown (not degradation). ‡ points to CDC25A-EGFP signal in *Chk1 WT* embryos that did not enter mitosis within 25 h after nuclear envelope assembly. The data are from one experiment with two *Chk1 WT* and four *Chk1 mKO* mice treated individually. The line of the *Chk1 WT* average is shown. The 2- to 3-cell anaphase timing is shown in Supplemental Fig. 4F. **E** Schematics of experimental design. **F, G** Line chart of fluorescence signal intensity of CDC25A-EGFP of individual *Chk1 WT* (**F**) and *Chk1 mKO* (**G**) 2-cell stage embryos scanned using time-lapse light-sheet microscopy from nuclear envelope assembly to nuclear envelope breakdown. Embryos were microinjected and treated with 2.5 μM RO3306 (CDK1 kinase inhibitor) as shown in **E**. Asterisk mark signal drop after nuclear envelope breakdown (not degradation). ‡ points to CDC25A-EGFP signal in *Chk1 WT* embryos that did not enter mitosis within 25 h after nuclear envelope assembly. The data are from one experiment with four *Chk1 WT* and four *Chk1 mKO* mice treated in two or four groups, respectively. The line of the *Chk1 WT* average is shown. The 2- to 3-cell anaphase timing is shown in Supplemental Fig. 4H.

In mammals, CDK1 is activated by cyclins and CDC25 phosphatases. CHK1 mediates degradation of all CDC25 phosphatases (Chen et al, 2003; Goto et al, 2019; Kramer et al, 2004; Löffler et al, 2006; Mailand et al, 2000; Peng et al, 1997; Sanchez et al, 1997; Santoni-Rugiu et al, 2000; Schmitt et al, 2006; Sorensen et al, 2003) but most importantly CDC25A (Goto et al, 2019; Mailand et al, 2000; Santoni-Rugiu et al, 2000; Sorensen et al, 2003; Zhao et al, 2002). CHK1 depletion/inhibition causes stable CDC25A levels (Goto et al, 2019; Lam et al, 2004; Sorensen et al, 2003; Tse et al, 2007) and premature mitotic entry (Mailand et al, 2002; Molinari et al, 2000). These observations suggest that CDC25A is the major CDC25 phosphatase during early embryo development and its stabilization after *Chk1* depletion could explain the precocious CDK1 activation and subsequent premature mitotic entry in *Chk1 mKO* embryos.

To ascertain if a high CDC25A level causes premature mitotic entry, we overexpressed CDC25A by microinjecting *Cdc25a-Egfp* and *H2b-mCherry* cRNAs into WT embryos and observed premature mitosis in both the first and the second embryonic division (**Supplemental Fig. S4C, D**). We then determined whether a high CDC25A level causes premature mitotic entry in 2-cell *Chk1 mKO* embryos using a similar microinjection approach and observed an accelerated first and the second mitosis in both *Chk1 WT* and *Chk1 mKO* embryos, which was more pronounced in *Chk1 mKO* embryos (**Supplemental Fig. S4E, F**). We also monitored CDC25A-EGFP intensity from nuclear envelope assembly in the 2-cell embryos until NEBD before the subsequent mitosis (**Fig. 3B**). *Chk1 WT* showed a peak in CDC25A-EGFP intensity at 1 to 1.5 h after nuclear envelope assembly in 2-cell embryos, with a subsequent signal decline until NEBD at 8.5 h or later (**Fig. 3C)**. The timing of CDC25A-EGFP degradation was consistent with CDC25A phosphatase being degraded in S phase of 2-cell embryos. *Chk1 mKO* embryos microinjected with *Cdc25a-Egfp* cRNA showed an increase in CDC25A-EGFP signal intensity followed by a rapid signal decline caused by CDC25A-EGFP signal dispersion into the cytoplasm during NEBD after mitotic entry (86%, n= 18) (**Fig. 3D**). These results suggest that a high CDC25A concentration causes premature mitotic entry in 2-cell *Chk1 mKO* embryos.

Because most *Cdc25a-Egfp* cRNA injected *Chk1 mKO* embryos divided too early to analyze the CDC25A-EGFP signal before NEBD, we delayed mitotic entry by partially inhibiting CDK1 with 2.5 µM RO3306. The zygotes were microinjected with cRNAs for *Cdc25a-Egfp* and *histone H2b-mCherry*, and then treated with RO3306 (**Fig. 3E**). This protocol successfully delayed the first and the second mitosis in *Chk1 WT* and *Chk1 mKO* embryos (**Supplemental Fig. S4G, H**) in compared to untreated *Chk1 WT* and *Chk1 mKO* embryos (**Supplemental Fig. S4E, F**). Next, we analyzed the CDC25A-EGFP signal in CDK1-inhibited 2-cell embryos. CDC25A-EGFP levels decreased more slowly in CDK1-inhibited *Chk1 WT* embryos (**Fig. 3F, Movie M3**) compared to non-inhibited *Chk1 WT* embryos (**Fig. 3C**). On the other hand, all CDK1-inhibited *Chk1 mKO* embryos showed CDC25A-EGFP signal stabilization (**Fig. 3G**) compared to non-inhibited *Chk1 mKO* embryos (**Fig. 3D**). We also noted that a subset of CDK1-inhibited *Chk1 mKO* embryos (referred as mKO1) showed WT-like timing of mitotic entry and slow decrease of CDC25A-EGFP levels similar to CDK1-inhibited *Chk1 WT* embryos (**Fig. 3G, Supplemental Fig. S4I, J; Movie M3**). On the other hand, a subset of CDK1-inhibited *Chk1 mKO* embryos (referred as mKO2) showed accelerated NEBD followed by CDC25A-EGFP dispersion into the cytoplasm (**Fig. 3G, Supplemental Fig. S4I, J; Movie M3**). These results imply that CHK1 regulates mitotic entry in 2-cell stage embryos by controlling CDC25A degradation.

### Maternally-expressed CDC25A regulates the first and second embryonic divisions

We next examined in more detail the role of CDC25A in early embryos. To determine whether CDC25A is required for the first embryonic division, we used maternal conditional knockout mice (*Cdc25a* F/F; Zp3-Cre;; *Cdc25a mKO*) or control mice (*Cdc25a* F/F;; *Cdc25a WT*), using *Zp3-Cre* to deplete the floxed *Cdc25a* allele specifically in oocytes. To produce *Cdc25a mKO* and *Cdc25a WT* embryos, *Cdc25a mKO* and *Cdc25a WT* females were mated with WT males.

We first determined *Cdc25a* mRNA abundance in *Cdc25a WT* and *Cdc25a mKO* in G2-phase zygotes (26 h post hCG administration) and G2-phase 2-cell embryos (42 h and 46 h post hCG administration). In *Cdc25a WT* embryos, *Cdc25a* mRNA abundance was high in zygotes in G2 at 26 h post hCG administration and substantially decreased in 2-cell embryos during the long G2 phase at 42 and 46 h after hCG administration. *Cdc25a* mRNA was virtually absent at all timepoints in *Cdc25a mKO* embryos (**Fig. 4A**). Thus *Cdc25a* transcript is absent (or very low abundance) during the long G2 phase in 2-cell embryos.

**Figure 4.**
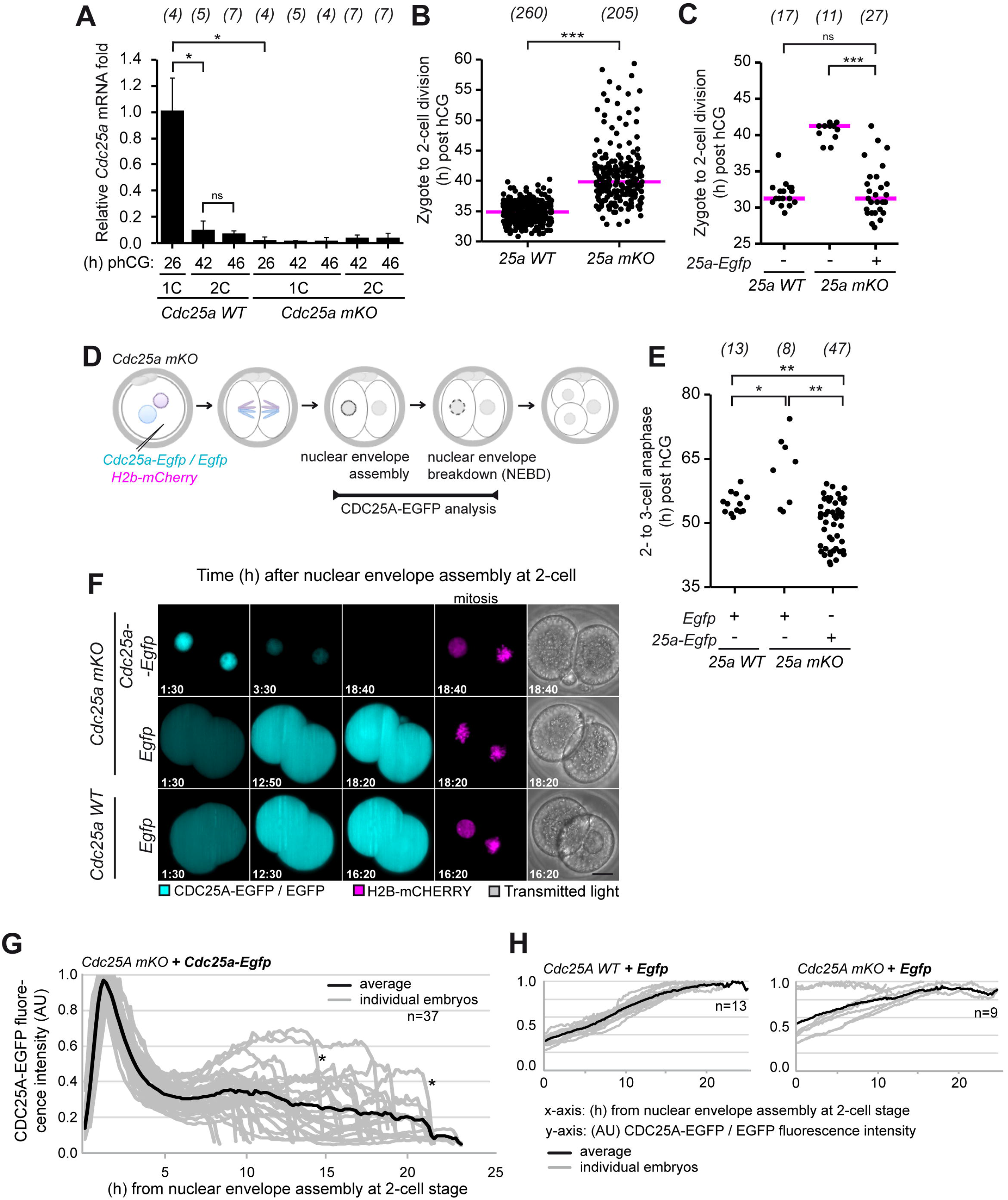
Maternally-expressed CDC25A regulates first and second embryonic division. **A** Bar graph of relative average *Cdc25a* mRNA folds from individual *Cdc25a WT* and *Cdc25a mKO* embryos from 26 to 46 h post hCG administration. The numbers in parentheses denote the number of embryos analyzed per group. 1C=1-cell, 2C=2-cell. The data are from one experiment with four *Cdc25a WT* and seven *Cdc25a mKO* mice treated individually. Data for each group are at least from 2-5 different mice. n.s. not significant, and *p<0.05 (Mann-Whitney U test, two-sided). **B** Dot-plot of the division time (h) from bright-field time-lapse imaging of zygote to 2-cell in *Cdc25a WT* and *Cdc25a mKO* embryos. The median is shown. The data were pooled from three independent experiments. ***p<0.001 (Mann-Whitney U test, two-sided). **C** Dot-plot of the division time (h) from bright-field time-lapse imaging from zygote to 2-cell in *Cdc25a WT*, non-injected *Cdc25a mKO* and *Cdc25a mKO* embryos microinjected with *Cdc25a-Egfp* cRNA. The median is shown. The data are from one experiment. Strain background: C57J/BL6 background with small part CD-1 or CD-1 background with small part C57J/BL6. n.s. not significant, and ***p<0.001 (Mann-Whitney U test, two-sided). **D** Schematics of experimental design. **E** Data from time-lapse light-sheet imaging (**E**) was used for quantification the division time from 2-cell to 3-cell in *Cdc25a WT* embryos expressing EGFP, and *Cdc25a* mKO embryos expressing EGFP or CDC25A-EGFP from microinjected cRNAs. The data were pooled from four independent experiments. *p<0.05, and **p<0.01 (Kolmogorov-Smirnov Test, two-sided). **F** Representative still images from time-lapse light-sheet imaging of preimplantation development of *Cdc25a WT* embryos expressing EGFP, and *Cdc25a* mKO embryos expressing EGFP or CDC25A-EGFP from microinjected cRNAs. Embryos were microinjected, as shown in **D**. Embryos expressing CDC25A-EGFP or EGFP (cyan) and H2B-mCHERRY (magenta, chromosomes) are shown. Scalebar 20 μm. Supplemental Movie 4 related to Fig. 4F: *Cdc25A mKO* + CDC25A-EGFP *Cdc25A mKO* + EGFP *Cdc25A WT* + EGFP **G** Line chart of fluorescence signal intensity of CDC25A-EGFP in *Cdc25a mKO* 2-cell stage embryos from nuclear envelope assembly to nuclear envelope breakdown. Asterisk mark signal drop after nuclear envelope breakdown (not degradation). **H** Line charts of fluorescence signal intensity of EGFP in *Cdc25a WT* or *Cdc25a mKO* 2-cell stage embryos from nuclear envelope assembly to nuclear envelope breakdown.

Because results described above suggested a role for CDC25A in regulating cell cycle progression, we examined the timing of cell division of *Cdc25a mKO* zygotes. Bright-field time-lapse imaging revealed a significant delay in cleavage of zygotes of *Cdc25a mKO* embryos (**Fig. 4B**) that was rescued by *Cdc25a-Egfp* cRNA microinjection (**Fig. 4C**). We then monitored the CDC25A-EGFP signal in *Cdc25a mKO* 2-cell embryos from nuclear envelope assembly to NEBD (**Fig. 4D, E, F; Movie M4**). Similarly, *Cdc25a mKO* embryos showed a delay in the second mitosis, and injection with *Cdc25a-Egfp* cRNA rescued the delay (**Fig. 4E**). Notably, the rescued *Cdc25a mKO* embryos showed a greater variability in the timing of the second embryonic mitosis, with a proportion showing the second embryonic mitosis earlier than the *Cdc25a WT* embryos (**Fig. 4E)**. A portion of the *Cdc25a mKO* embryos showed WT-like timing of the second anaphase, which could have reflected a CDC25A-EGFP concentration similar to that of wild-type CDC25A in WT embryos. In these embryos, the CDC25A-EGFP signal rapidly increased with a peak corresponding to the time of the G1/S transition followed by CDC25A-EGFP degradation corresponding to the time of S phase progression (**Fig. 4G**). Most embryos then showed a slight CDC25A-EGFP signal increase around 6-10 h post-nuclear envelope assembly, followed by a rapid signal decrease after NEBD during mitotic entry or by a slight signal decrease. The signal from EGFP alone increased gradually from the nuclear envelope assembly until NEBD suggesting that changes in CDC25A-EGFP intensity is not due to selective translation or degradation of the EGFP moiety of the fusion protein (**Fig. 4G, H**). Thus, in 2-cell embryos, CDC25A is degraded during S phase progression and remains lower thereafter. Taken together, these results indicate that CDC25A is essential for regulating cell cycle progression of zygotes and 2-cell embryos.

### Maternal CHK1 regulates chromosome segregation, and its absence causes genome fragmentation

CHK1 is a critical DDR kinase (Takai et al, 2000). Its depletion causes chromosome instability in 8-cell embryos (Ruth et al, 2021) and DNA damage, premature mitosis, chromosome fragmentation, and micronuclei formation in cell lines (Dandoulaki et al, 2018; Durkin et al, 2006; Hoffelder et al, 2004; Niida et al, 2005; Peddibhotla et al, 2009). Therefore, we next examined chromosome stability in *Chk1 mKO* embryos using long-term time-lapse imaging on a light-sheet microscope and visualized chromosomes (**Fig. 5A**).

**Figure 5.**
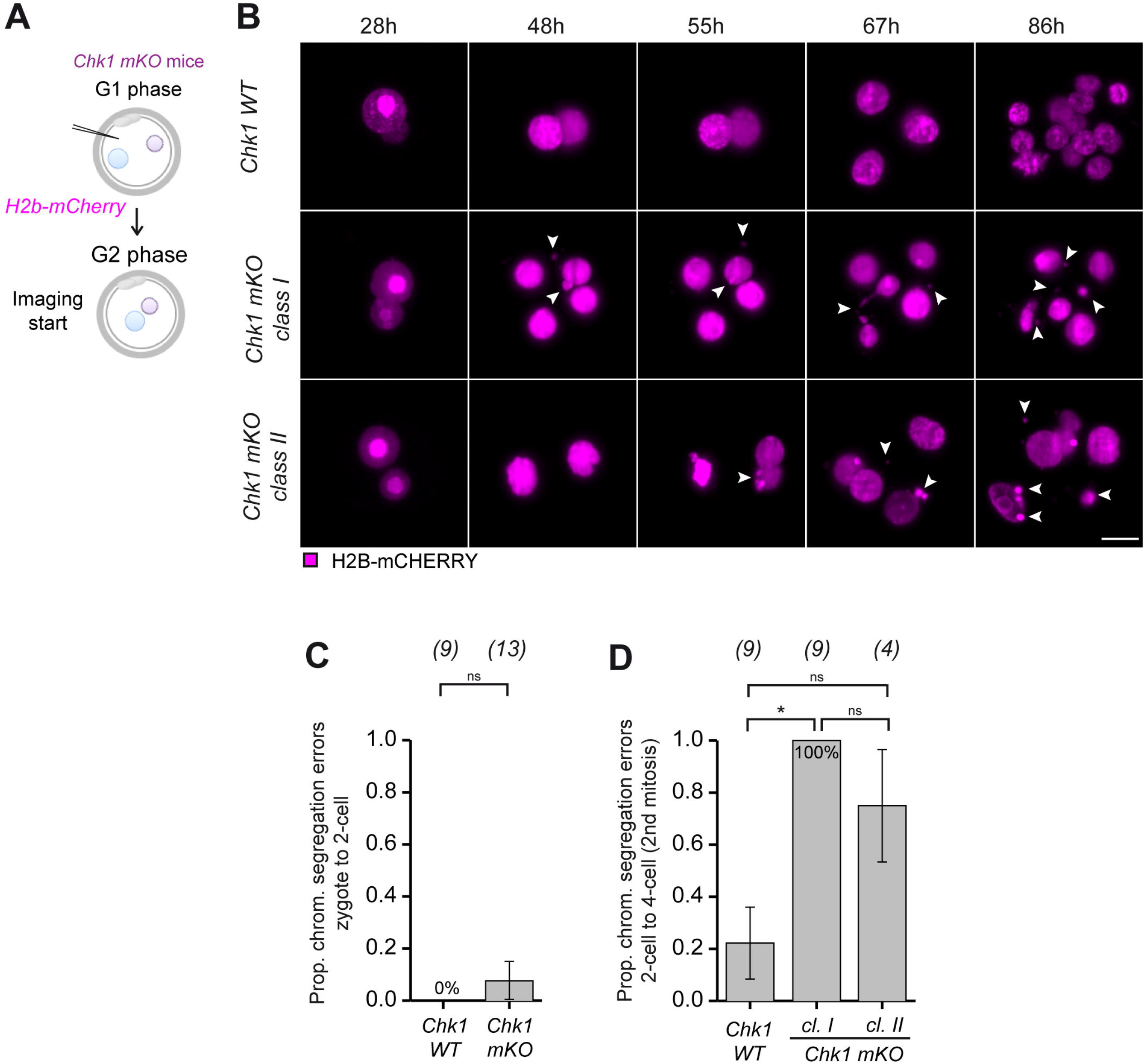
Massive genome fragmentation in *Chk1 mKO* embryos is caused by chromosome segregation errors during the second mitosis. **A** Schematic of the experimental procedure. **B** Representative still images from time-lapse light-sheet imaging of *Chk1 WT, Chk1 mKO class I*, and *Chk1 mKO class II* embryos from 28 to 86 h post hCG administration. Embryos were microinjected, as shown in **A**. Embryos expressing H2B-mCHERRY (magenta, chromosomes) are shown. Timepoints are relative to time post hCG administration (h). Arrowheads mark genome fragmentation. Scalebar 20 μm. Movie M5 related to Fig. 5B: Chk1 WT Chk1 mKO class I Chk1 mKO class II **C, D** Data from time-lapse light-sheet imaging (**B**) was used to quantify the percentage of filmed embryos with chromosome segregation errors in the first embryonic division from zygote to 2-cell stage (**C**) and in the second embryonic division from 2-cell to 3-cell stage and 3-cell stage to 4-cell stage (**D**). Chromosome segregation errors were counted if observed in at least one of these divisions. The numbers in brackets denote the number of analyzed embryos per genotype. The data were pooled from two independent experiments. Error bars denote the standard error of proportion. n.s. not significant, and *p<0.05 (Fisher’s Exact Test).

In the first division (zygote to 2-cell stage), anaphase was accelerated by 1 h in *Chk1 mKO* embryos compared to *Chk1 WT* (**Supplemental Fig. S5A, D; Movie M5**). Also, *Chk1 mKO* embryos showed a slight but non-significant increase in chromosome segregation errors during the first anaphase (**Fig. 5B, C; Movie M5**). In individual *Chk1 mKO* embryos, only mild segregation errors were observed during the first anaphase (**Supplemental Fig. S5D; Movie M5**). The acceleration of anaphase in *Chk1 mKO* embryos was more pronounced at the 2- to 3-cell division (**Supplemental Fig. S5B**), and by the 4-cell stage, nearly 80% displayed chromosome segregation errors compared to 20% of control embryos (and more pronounced in *Chk1 mKO class I* than *Chk1 mKO class II* embryos (**Fig. 5D, Supplemental Fig. S5C, E**). The chromosome segregation errors were lagging chromosomes, anaphase bridges, and genome fragmentation (**Supplemental Fig. S5C, F; Movie M6**). We also observed these segregation errors in *Chk1 mKO* embryos that showed WT timing of the initial embryonic divisions. Thus, CHK1 is essential for genome integrity during the initial embryonic divisions after fertilization.

### Genome fragmentation after *Chk1* depletion is not transcription-dependent

Next, we sought to understand the basis for the chromosome segregation errors after *Chk1* depletion. CHK1 is a component of the replication stress response, which can be triggered by transcription-replication collisions (Barroso et al, 2019; Blythe & Wieschaus 2015; Lam et al, 2020; Sollier & Cimprich 2015). Transcriptional activity in mouse 2-cell embryos is high because of the major phase of genome activation (Aoki et al, 1997). Therefore, we asked whether preventing zygotic genome activation would suppress chromosome segregation errors in *Chk1 mKO* embryos (**Fig. 5D, Supplemental Fig. S2G**).

We inhibited RNA polymerase II and III by culturing zygotes in α-amanitin (24 µg/ml) from 26 h after hCG administration (**Fig. 6A**). At this time, the majority of embryos have completed the first S phase (Qiu et al, 2003; Rambhatla & Latham 1995; Zeng & Schultz 2005) but have not yet initiated major zygotic genome activation (Aoki et al, 1997; Hamatani et al, 2004). α-amanitin treatment did not rescue the premature division of *Chk1 mKO* embryos (**Supplemental Fig. S6A**). Whereas α-amanitin treatment led to a robust 2-cell stage block in *Chk1 WT* embryos even at 68 h (**Fig. 6B, C**), a robust 2-cell stage block depended on *Chk1*, because 62% of the α-amanitin-treated *Chk1 mKO* embryos progressed to the 3 or 4 cell stage at 68 h before arresting (**Fig. 6B, C**). These embryos remained in the 2- or 3-/4-cell stage until 114 h post hCG stimulation, when 69% of DMSO-treated *Chk1 WT* developed into blastocyst stage (**Supplemental Fig. S6B, C**). Increased genome fragmentation, as assayed by DAPI staining, was observed in 4-cell *Chk1 mKO* embryos cultured in the presence of α-amanitin (24 µg/ml) as well as DMSO (**Fig. 6D**) from 26 h-68 h post hCG administration (**Fig. 6E, F**). Thus, when major genome activation is inhibited in *Chk1 mKO*, cleavage arrest depends on CHK1 activity, but genome fragmentation is not transcription-dependent.

**Figure 6.**
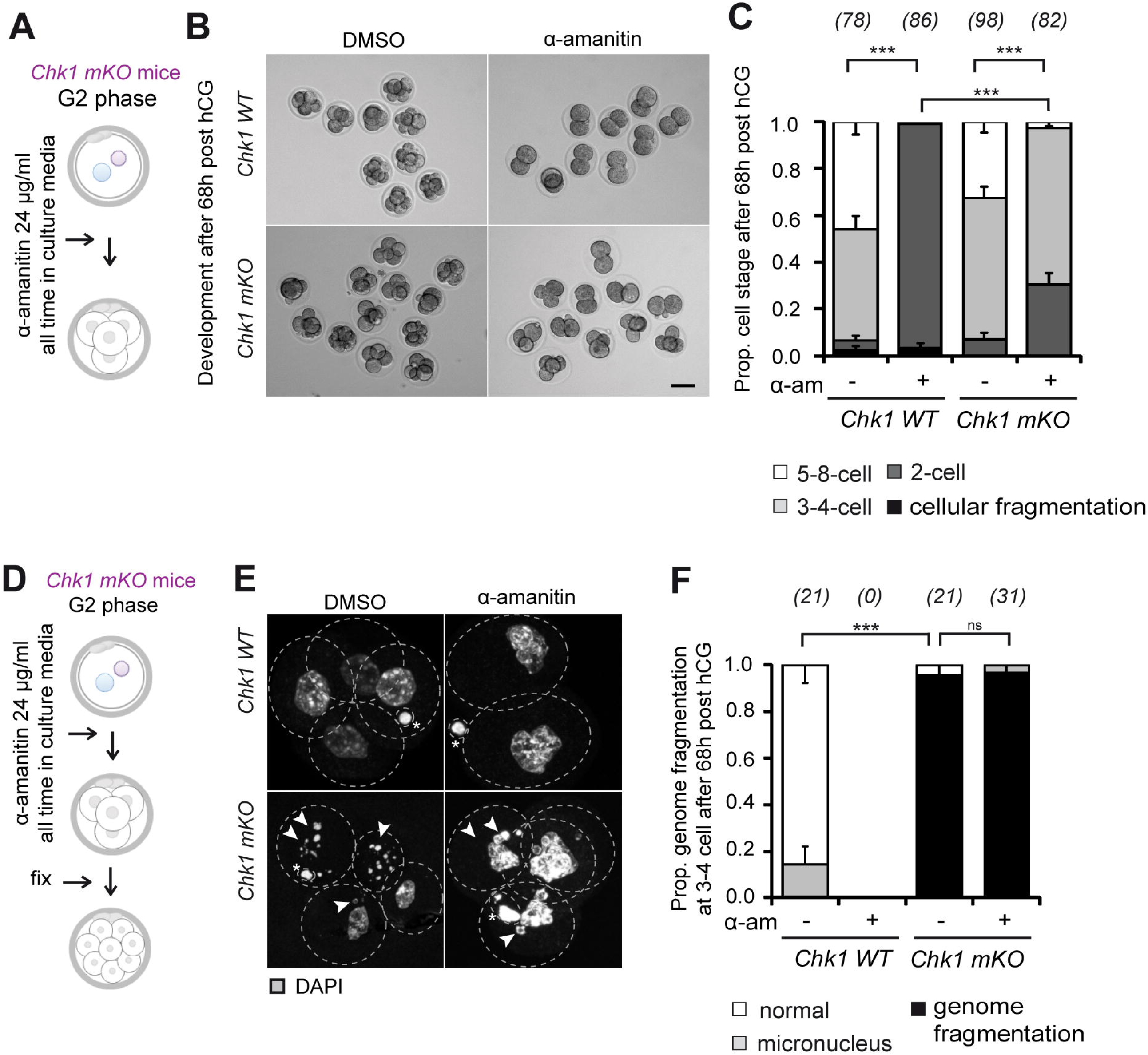
DNA damage and chromosome segregation errors in *Chk1 mKO* embryos are not transcription-dependent. **A** Schematic depiction of the experimental procedure. **B** Representative images from developmental analysis of *Chk1 WT* and *Chk1 mKO* embryos treated with DMSO or α-amanitin (24 µg/ml). Embryos were scanned at 68h post-hCG administration. Strain background: CD-1 with small part C57J/BL6 and C57J/BL6 with small part CD-1. Scalebar 60 μm. **C** Proportion of developmental stages of embryos (as shown in **B**) at 68 h post hCG administration. The numbers in parentheses denote the number of analyzed embryos per group. The data were pooled from three independent experiments. Error bars denote standard error of proportion. ***p<0.001 (Cochran-Armitage Trend Test, two-sided). **D** Schematic depiction of the experimental procedure. **E** Representative images from genome integrity analysis of *Chk1 WT* and *Chk1 mKO* embryos treated with DMSO or α-amanitin (24 µg/ml). The embryos were scanned at 68 h post-hCG administration. Embryos stained with DAPI (gray, chromatin) are shown. A dashed line indicates blastomeres outlines as seen in a bright field. Arrowheads mark genome fragmentation. Asterisk marks polar bodies. Strain background: CD-1 with small part C57J/BL6 and C57J/BL6 with small part CD-1. **F** Proportion of genome fragmentation of embryos (as shown in **E**). Only 3-4 cell stages were analyzed. Note that all *Chk1 WT* embryos were at the 2-cell stage. The data were pooled from two independent experiments. Error bars denote standard error of proportion. n.s. not significant, and ***p<0.001 (Cochran-Armitage Trend Test, two-sided).

### The long G2 phase in 2-cell embryos is necessary for genome integrity

We suspected that the cause for the high incidence chromosome segregation errors after *Chk1* depletion during the second division was due to premature mitosis due to a shortened G2 phase. Accordingly we forced *H2b-Egfp* transgenic embryos into premature mitosis by WEE/MYT pharmacological inhibition (Ferencova et al, 2022) at 44h post hCG administration (**Fig. 7A**). At this time, 92% of WT embryos were in G2, because we did not detect DNA replication following an EdU pulse (**Supplemental Fig. S3F, G**). WEE/MYT-inhibited 2-cell embryos entered mitosis 7 h earlier than controls (**Fig. 7B**), and was accompanied by segregation errors in 66% of the embryos as well as an extended M phase (1.8 h compared to 0.8 h in WT) (**Fig. 7C, D**). Thus, the long G2 phase in 2-cell embryos—and maintaining that long G2 phase—is essential for genome integrity.

**Figure 7.**
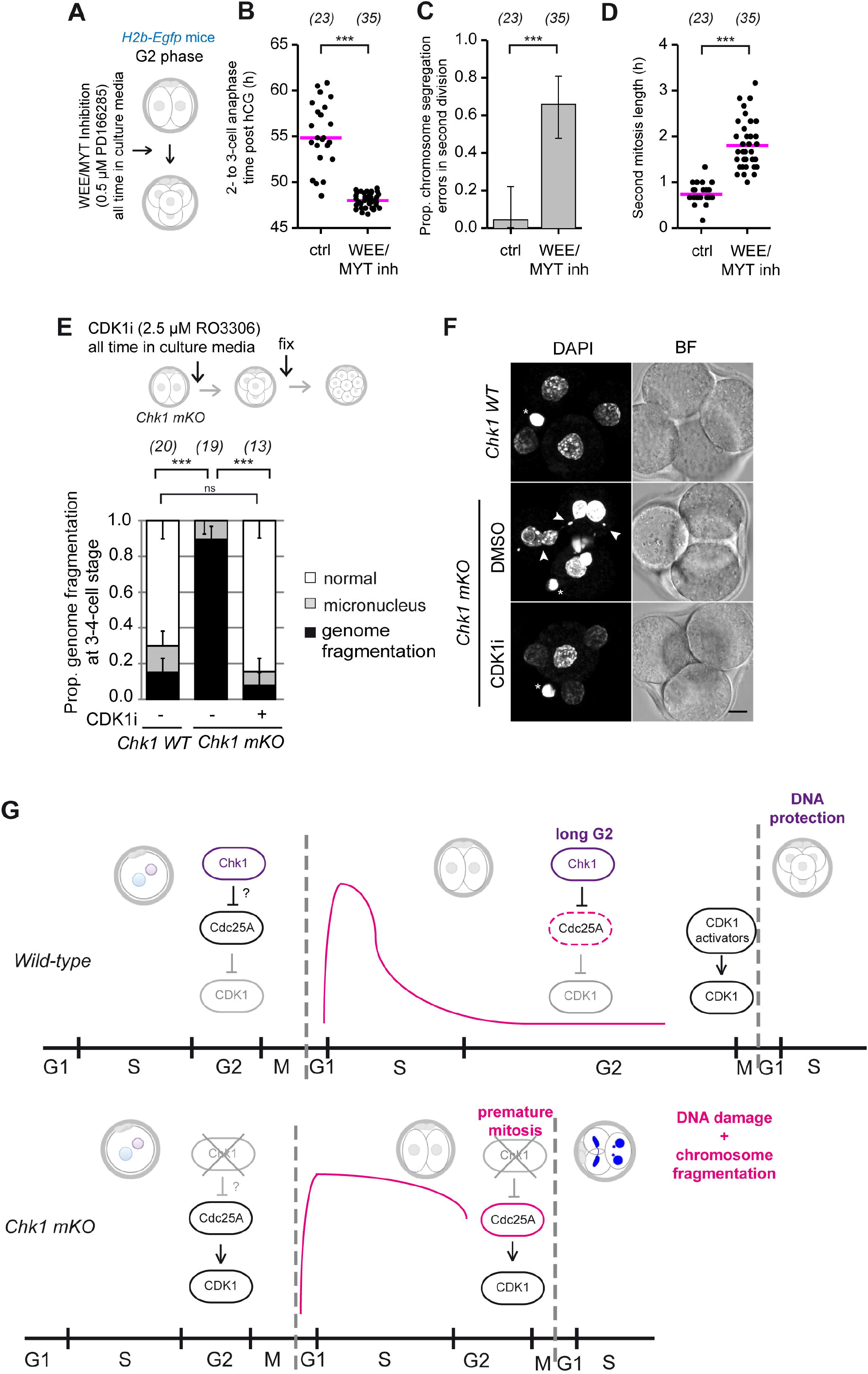
CHK1 is necessary for genome integrity during long G2 in 2-cell stage mouse embryos. **A** Schematic depiction of the experimental procedure. **B, D** Data from time-lapse light-sheet imaging of embryos from *H2b-Egfp* transgenic mice after WEE/MYT kinases inhibition (as shown in **A**) at 44 h post hCG administration (G2 phase) was used to quantify the initiation of anaphase (h) from 2-cell to 3-cell stage embryos (**B**), and length of mitosis (h) in 2-cell to 3-cell stage embryos (**D**). The numbers in parentheses denote the number of analyzed embryos per group. The data were pooled from two independent experiments. The median is shown. ***p<0.001 (Aspin-Welch Unequal – Variance T-Test). **C** Data from time-lapse light-sheet imaging of embryos from *H2b-Egfp* transgenic mice after WEE/MYT kinases inhibition (as shown in **A**) at 44 h post hCG administration (G2 phase) was used to quantify the percentage of filmed embryos with chromosome segregation errors. Error bars denote the 95% confidence interval. ***p<0.001 (Fisher’s Exact Test, two-sided). **E** Genome integrity analysis of *Chk1 WT*, DMSO-treated *Chk1 mKO*, and RO3306-treated *Chk1 mKO* (CDK1 inhibition) within 3 h after 2-cell division. The embryos were scored at 60 h post hCG administration, and only 3-4 cell-stage embryos were analyzed. The data are from one experiment with four groups of different mice of *Chk1 mKO* and one group of *Chk1 WT*. Error bars denote standard error of proportion. n.s. not significant, and ***p<0.001 (Cochran-Armitage Trend Test, two-sided). **F** Representative images from genome integrity analysis of embryos (**E**). The embryos were scanned at 60 h post hCG administration. Embryos stained with DAPI (gray, chromatin) are shown. Arrowheads mark genome fragmentation. Asterisk mark polar bodies. BF bright-field. Scalebar 20 µm. **G** Model of cell cycle regulation and genome integrity protection by CHK1 in early mouse embryos. In wild-type mouse embryous, CHK1 restrains CDK1 activity by promoting CDC25A degradation to regulate cell cycle progression and protect genome integrity in 2-cell embryos. CDK1 is activated by its newly transcribed activators after genome activation.

The G2/M transition is regulated by CDK1 kinase. Therefore, we asked whether the DNA damage and chromosome segregation errors in *Chk1 mKO* embryos is a consequence of premature CDK1 activation. To this end, we partially inhibited CDK1 in 2-cell embryos with 2.5 µM RO-3306 and analyzed DNA integrity in 4-cell embryos. Partially CDK1-inhibited *Chk1 mKO* embryos showed reduced genome fragmentation compared to DMSO-treated *Chk1 mKO* embryos (**Fig. 7E**). The incidence of genome fragmentation in the *Chk1 mKO* embryos exposed to partial CDK1 inhibition was similar to that in *Chk1 WT* embryos (**Fig. 7E, F**). Taken together, these findings show that partial inhibition of CDK1 rescues both M phase timing and genome fragmentation in 2-cell *Chk1 mKO*. We infer that CHK1 restrains CDK1 activity by promoting CDC25A degradation to regulate cell cycle progression and protect genome integrity in 2-cell embryos (**Fig. 7G**).

## DICUSSION

Cell cycle progression during early development is accompanied by shortening and/or lengthening of G1 or G2 compared to somatic cells. For *Drosophila, Xenopus*, zebrafish, and mammals, fertilization is followed by a long G1 (Hörmanseder et al, 2013), in contrast to a short or absent G1 before the major phase of genome activation **(Fig. 2)** (Artus & Cohen-Tannoudji 2008; Palmer & Kaldis 2016). Genome activation depends on lengthening G1 or G2 (Artus & Cohen-Tannoudji 2008; Braude 1979; Flach et al, 1982; Foe & Alberts 1983; Liu & Grosshans 2017; Newport & Kirschner 1982; Zamir et al, 1997).

Regulation of the long G2 phase in mammalian embryos is poorly understood and it was thought that the G2/M depended on transcription during genome activation (Braude 1979; Flach et al, 1982). Results reported here show that the G2/M transition in mouse 2-cell embryos is *Chk1*-dependent where CHK1 prevents premature mitosis by restraining CDK1 activity by inducing CDC25A degradation. Thus, we propose that the long G2 phase is normally maintained by the absence of CDC25A until CDK1 is activated by its newly transcribed activators after genome activation (**Fig. 7G**). In support of this model is that maternal *Chk1* depletion caused decrease in percentage of G2 zygotes (**Supplemental Fig. S2D**) and shortens the length of G2 in 2-cell embryos (**Fig. 2**), with a premature mitosis following in both cases (**Fig. 1**); premature mitotic entry in 2-cell *Chk1 mKO* embryos results from CDC25A protein stabilization (**Fig. 3D, G**) and CDC25A overexpression induces premature mitosis (**Supplemental Fig. S4C, D**). Generation of newly transcribed CDK1 activators is consistent with an increase in transcript abundance of mitotic activators such as CDK1, CDK2, and cyclins D1-3, E1, A2, and CDC25C from the zygote to the 2-cell stage (Abe et al, 2018). The low levels of *Cdc25a* mRNA in 2-cell embryos may therefore become rate-limiting for cell cycle progression, which is delayed until e.g., *Cdc25c* mRNA levels increase of the major genome activation (Park et al, 2013; Xie et al, 2010; Zeng et al, 2004). In support of this model also is that CDC25A-EGFP is degraded during the time similar to S-phase timing in WT 2-cell embryos (**Fig. 3C)** and CDK1 inhibition rescues the G2 duration and the timing of M phase in *Chk1 mKO* embryos (**Fig. 3A**). This model is consistent with others. *Chk1* depletion causes aberrant embryonic development in mouse (Liu et al, 2000; Takai et al, 2000). CHK1 inhibition causes premature mitosis in both mouse (Ju et al, 2020) and human (Palmerola et al, 2022) zygotes. CHK1 kinase activity itself is necessary for early embryos (Muralidharan et al, 2020). It is also consistent with more other studies in mice and (Fishler et al, 2010; Lam et al, 2004; Niida et al, 2005; Peddibhotla et al, 2009; Ruth et al, 2021), humans (Branigan et al, 2021; Durkin et al, 2006; Niida et al, 2005; Syljuåsen et al, 2005) and other species (Zachos et al, 2007, 2003, 2005).

There are evolutionary conserved features and differences in CHK1’s role in regulating cell cycle length in embryonic development. In non-mammalian models, genome activation is preceded by 9-14 rapid cell cycle divisions (Blythe & Wieschaus 2015; Collart et al, 2013; Dalle Nogare et al, 2009), during which ATR-CHK1 is prevented from recognizing replication stress (Blythe & Wieschaus 2015; Holway et al, 2006; Shimuta et al, 2002; Sibon et al, 1997; Zhang et al, 2014). At the time of genome activation, CHK1 causes CDC25A degradation in both zebrafish (Dalle Nogare et al, 2009; Zhang et al, 2014), *Drosophila* (Deneke et al, 2016) and *Xenopus* (Petrus et al, 2004; Shimuta et al, 2002), as in mouse reported here (**Fig. 3**). Whether CHK1 in mammals regulates other CDKs and CDC25s during genome activation during the maternal-to-zygotic transition warrants further investigation, noting that CHK1 overexpression in *Xenopus* egg extracts causes CDC25A degradation without affecting CDC25C levels (Hartley et al, 1996; Petrus et al, 2004) and inhibiting CDC25A and CDK1 but not CDK2 causes shortening of the long G2 during genome activation in zebrafish (Dalle Nogare et al, 2009).

Our results are consistent with a step-wise increasing activation of CDK1 in early embryos such that by reaching certain thresholds during cell cycle progression specific cell cycle transitions/phases (G1/S transition, S phase progression, G2 length, and mitotic entry) occur (**Fig. 2**) (Branigan et al, 2021; Lemmens et al, 2018; Niida et al, 2005; Suski et al, 2022). This model implies that for a premature mitotic entry from early or middle S phase, CDK1 activity needs to be increased more than for a premature mitotic entry from the late S or G2 phase (Pomerening et al, 2003; Sha et al, 2003). Consistent with this proposal is that CHK1 depletion/inhibition or CDC25A-EGFP overexpression alone causes premature mitotic entry only from the late S or G2 phase (**Fig. 2**; **Supplemental Fig. S4F**) (Branigan et al, 2021; Lemmens et al, 2018; Niida et al, 2005). For premature mitotic entry from the early or middle S phase 1) a concomitant *Chk1* depletion and CDC25A-EGFP co-expression (**Fig. 3D; Supplemental Fig. S4F**), or 2) separate WEE/MYT kinase inhibition (**Supplemental Fig. S3C, Fig. 7B**) are required.

The extent to which CHK1 regulates replication completion is not fully understood. Although *Chk1 mKO class I* embryos exhibit premature mitotic entry after finishing the main period of S phase progression (**Fig. 2**), this observation does not necessarily mean they completed replication. The transition to G2 is set by a low threshold of origin firing (Zonderland et al, 2022), but in some cases, replication can continue during G2 phase (Mocanu et al, 2022; Palmerola et al, 2022). This finding is consistent with other studies (Branigan et al, 2021; Lam et al, 2004), where *Chk1* depletion or inhibition causes premature mitosis in cells with only mildly, but not severely, under-replicated DNA.

CHK1 is activated after replication stress in somatic cells and early embryos (González Besteiro & Gottifredi 2015; Lam et al, 2004; Lebrec et al, 2022; Lemmens et al, 2018; Michelena et al, 2019; Moiseeva et al, 2019; Muralidharan et al, 2020; Palmerola et al, 2022; Syljuåsen et al, 2005; Zachos et al, 2005). Replication stress signalling is usually orchestrated by ATR kinase (Eykelenboom et al, 2013; Saldivar et al, 2018). However, the role of ATR in early embryos does not seem as prominent as CHK1. *Atr* germ-line KO embryos show better development than *Chk1* germ-line KO embryos (Brown & Baltimore 2000; Liu et al, 2000; Takai et al, 2000).

Disrupting CHK1-CDC25A-CDK1 regulation results in DNA damage (**Supplemental Fig. S2G**), chromosome segregation errors (**Fig. 5D**), aneuploidies, and infertility (Ruth et al, 2021). These findings are consistent with studies in mice (Fishler et al, 2010; Lam et al, 2004; Niida et al, 2005; Peddibhotla et al, 2009; Ruth et al, 2021), humans (Branigan et al, 2021; Durkin et al, 2006; Niida et al, 2005; Syljuåsen et al, 2005) and other species (Zachos et al, 2007, 2003, 2005). We observed, however, that the DNA damage and chromosome segregation errors after *Chk1* depletion occur only in 2-cell embryos (**Supplemental Fig. S2G, Fig. 5D**) and not in zygotes (**Supplemental Fig. S2A, Fig. 5C**). This finding contrasts with human zygotes, where CHK1 inhibition causes both DNA damage and segregation errors in 9 out of 9 embryos, noting that development of human embryos is more error-prone (Cavazza et al, 2021; preprint: Ford et al, 2020; Palmerola et al, 2022; Vanneste et al, 2009) than inbred mice (Bolton et al, 2016; Mashiko et al, 2020). These observations suggest an evolutionary difference (Svoboda 2018), where zygotes from inbred mice are more efficient in protecting their DNA integrity.

Infertility rates worldwide are increasing (Carson & Kallen 2021; Gruhn & Hoffmann 2022). About 50% of human embryos do not develop to the blastocyst stage due to a high incidence of aneuploidy (preprint: McCoy et al, 2022). Also, almost 95% of human arrested embryos show chromosome segregation errors (preprint: McCoy et al, 2022), and even in blastocysts, 20% show mosaicism when disaggregated (Capalbo et al, 2021) or 30% when a biopsy is analyzed (preprint: McCoy et al, 2022). There is increasing evidence that some of these aneuploidies are of mitotic (embryonic) origin (preprint: McCoy et al, 2022; Vanneste et al, 2009). Results reported here using a mouse model implicate CHK1-CDC25A-CDK1 regulation as another locus whose dysregulation could contribute to human infertility.

## METHODS

### Animal use and *Chk1 mKO* generation

Mice were handled and experiments conducted accordingly to the guidelines of the Expert Committee for the Approval of Projects of Experiments on Animals of the Academy of Sciences of the Czech Republic (no. 43-2015). Mice were housed at the Institute of Animal Physiology and Genetics of the Czech Academy of Sciences in Libechov, Czech Republic. Mice were housed under conditions of a 12 h day-night cycle at 22 °C (±2 °C), and had access to food and water ad libitum.

The *Chk1 mKO* embryos were created as in (Ruth et al, 2021) by crossing mice carrying a Cre-recombinase expressed from the oocyte-specific *Zp3* promoter (Lewandoski et al, 1997) to mice with floxed *Chk1* alleles (Lam et al, 2004). All experiments were conducted by using littermates. The mice were on CD-1 background with a small part of C57J/BL6 if not noted otherwise in Figure legends.

The *Cdc25a mKO* embryos were created by crossing mice carrying a Cre-recombinase expressed from the oocyte-specific *Zp3* promoter (Lewandoski et al, 1997) to mice with floxed *Cdc25a* alleles (Lee et al, 2009). All experiments were conducted by using littermates. The mice were on C57J/BL6 background with a small contribution of CD-1 if not noted otherwise in Figure legends.

*H2b-Egfp* mice were on CD-1 background (Mayer et al, 2016).

WT strain of mice was either CD-1 (ICR) or C57J/BL6.

### Genotyping

Animals were genotyped two times, initially upon weaning and again after experimental procedures were carried out.

*Chk1 mKO* genotyping was performed by PCR analysis using the following primers for *mKO Chk1(Zp3-Cre):* CHK1F1 (5’-ACC TGC CCG CAA CTC CCT TTC-3’), CHK1R1(5’-CCA TGA CTC CAA GCA CAG CGA-3’), Cre_low (5’-TAT TCG GAT CAT CAG CTA-3’), Cre_up (5’-

GGT GGG AGA ATG TTA ATC-3’). The sizes of products were 318 bp for WT and 380 bp for lox/lox transgene. The size of the *Zp3-*Cre transgene was 139 bp. The *Taq* 2X Master Mix was used for genotyping (NEB, # M0270L).

*Cdc25a mKO* genotyping was performed by PCR analysis using the following primers for *Cdc25a mKO (Zp3-Cre):* oIMR9611: (CAG AGC CTG AAG TCC TGT GAA GG), oIMR9612: (CTG GGT AGT GTA GTT CCT ACA GCG), Cre_low (5’-TAT TCG GAT CAT CAG CTA-3’), Cre_up (5’-GGT GGG AGA ATG TTA ATC-3’). The sizes of products were 222 bp for WT and 270 bp for lox/lox transgene. The size of the *Zp3-*Cre transgene was 139 bp. The *Taq* 2X Master Mix was used for genotyping (NEB, # M0270L).

### Superovulation and embryo handling

Mice 3-6 months old were stimulated with 5 IU of PMSG (HOR-272, ProSpec-Tany TechnoGene), and after 44 h with 5 IU of hCG (Ovitrelle, Merck) and then mated with a WT CD-1 male. The females were sacrificed by cervical dislocation after 18 h in accordance with the guidelines of ethics committee.

The oviducts were placed in M2 medium (M7167, Merck), and the ovulated MII oocytes and zygotes were collected from teared ampullae. The cumulus mass containing the MII oocytes and zygotes was transferred to a drop of M2 medium containing 300 µg/ml hyaluronidase (H4272, Merck) to release the cumulus cells. The MII oocytes and zygotes were cultured in EmbryoMax KSOM medium (MR-106-D, Merck) at 37 °C with 5% CO_2_ in an atmosphere of air. For live-cell imaging, embryos were cultured in global® (LGGG-020, Origio) supplied with 1 mg/ml BSA. Only zygotes with visible pronuclei were used for experiments.

The temporal information about embryo development is given in (h) post hCG administration, as this is the last exact time-point in mating with males.

### Oocyte handling

For oocyte collection, mice were stimulated with 5 IU of PMSG (HOR-272, ProSpec-Tany TechnoGene) and the germinal vesicle (GV)-stage oocytes were collected in M2 medium supplied with 2.5 µM milrinone (M4659, Merck) to prevent resumption of meiosis. The oocytes were cultured in minimum essential medium (MEM, Merck, M4655) containing 1.14 mM sodium pyruvate (Merck, P4562), 4 mg/ml bovine serum albumin (BSA, Merck, A3311), penicillin-streptomycin (75 U/ml-60 μg/ml, Merck, P7794, S1277) and incubated at 37°C in 5% CO_2_ in an atmosphere of air.

### Inhibitors

All used inhibitors were diluted in DMSO (Merck, D2650). The concentrations used were as follows: CHK1 inhibitor (CHIR-124) 250 µM, CDK1 inhibitor (RO-3306) 2.5 µM, WEE/MYT inhibitor (PD0166285) 0.5 µM (Ferencova et al, 2022).

### Plasmid preparation

The pCMV-Sport6-Chk1 plasmid containing mouse *Chk1* cDNA (NM_007691.5, from nucleotide 117 to 1541) and the pBSK-EGFP plasmid were kindly provided by Martin Anger, IAPG CAS, Libechov and CEITEC, Brno, Czech Republic. *Chk1* coding sequence was cloned into pBSK-EGFP using SpeI restriction sites to create pBSK-EGFP-Chk1 plasmid. To create pYX-Chk1 plasmid, EGFP sequence was replaced by *Chk1* coding sequence in pYX-EGFP (Blengini et al, 2021) using SpeI restriction sites.

To create a *Cdt1* sensor suitable for mouse oocytes and embryos, we modified an mCherry-hCdt1(1-100)Cy(-)pcDNA3 plasmid (RIKEN BRC DNA Bank) (Sakaue-Sawano et al, 2017). We performed codon optimization using IDT Codon Optimization Tool (Integrated DNA Technologies). The modified *mCdt1* sequence was synthesized (Integrated DNA Technologies) and cloned into pYX-EYFP (Blengini et al, 2021) using SpeI restriction sites to create pYX-EYFP-*mCdt1* plasmid for transcription of *mCdt1*-EYFP sensor.

Other plasmids for *in vitro* transcription were already described – pBSK-H2B-mCherry (Kitajima et al, 2011), pBSK-EGFP-Cdc25A (Solc et al, 2008), pXY-EGFP (Blengini et al, 2021; Saskova et al, 2008).

### cRNA production and microinjection

For cRNA preparation, plasmids were linearized by AscI (pBSK vector) or SfiI (pYX vector) and purified by NucleoSpin Gel and PCR Clean-up kit (Macherey-Nagel, #740609). cRNA was produced by in *vitro* transcription using mMessage mMachine™ T3 kit (Ambion, #1348). *Chk1-Egfp, Chk1, Egfp*, and *Cdc25a-Egfp* cRNAs were polyadenylated using the Poly(A) Tailing Kit (Ambion, #AM1350) according to manufacturer’s protocols. cRNAs were purified using RNeasy Mini Kit (Qiagen, #74104) and stored at -80°C. The oocytes or embryos were microinjected with ∼10 pl of an cRNA solution (35 ng/μl *H2b-mCherry*, 10 ng/ul *Egfp-Chk1* or *Egfp* alone for *Chk1 mKO* rescue in **Fig. 1G**, 300 ng/μl *Chk1* alone for immunoblotting, 50, 75 or 100 ng/μl *Egfp-Cdc25a* and 50 ng/ul *Egfp* alone.

### Single-oocyte/embryo real-time qPCR

Expression level of *Cdc25a* mRNA was measured by RT-qPCR using FastLane Cell SYBR Green kit (Qiagen, #216213) using CFX96 Touch Real-Time PCR Detection System, Biorad. Single oocytes or embryos were directly used as a substrate for single-step RT-qPCR without mRNA purification. External *Gfp* mRNA (0.04 pg *Gfp* mRNA per oocyte/embryo) was added to lysed oocytes or embryos to serve as a control for RNA quantification and normalization of RT-qPCR data. The following gene-specific primers were used: *Cdc25a* primers located at 5′ end of mRNA (AATAACAGCAGTCTACAG and TTCAAGGTTTTCTTTACTG); *Cdc25a* primers located at 3′ end of mRNA (AGAACCCTATTGTGCCTACTG and TACTCATTGCCGAGCCTATC); *Gfp* primers (TTCAAGATCCGCCACAAC and GACTGGGTGCTCAGGTAG). The following single-step RT-qPCR protocols were used (modified from (Solc et al, 2008): *Cdc25a* 5′ end: 1. 50°C 30 min, 2. 95°C 15 min, 3. 94°C 15 sec, 4. 52°C 30 sec, 5. 72°C 30 sec, 6. 72°C 3 sec, 7. Plate reading, 8. go to step 3 for 50 additional cycles; *Cdc25a* 3′ end: 1. 50°C 30 min, 2. 95°C 15 min, 3. 94°C 15 sec, 4. 57°C 30 sec, 5. 72°C 30 sec, 6. 72°C 3 sec, 7. plate reading, 8. go to step 3 for 50 additional cycles; *Gfp*: 1. 50°C 30 min, 2. 95°C 15 min, 3. 94°C 15 sec, 4. 50°C 30 sec, 5. 72°C 30 sec, 6. 72°C 3 sec, 7. plate reading, 8. go to step 3 for 50 additional cycles. Following quantification of the PCR reaction product, expected size of product was verified by electrophoresis using 1% agarose gel. The product sizes were following; *Cdc25a* 5′ end – 99 bp, ; *Cdc25a* 3′ end – 124 bp, *Gfp* – 122 bp. Raw data without smoothening and with a global minimum baseline correction were imported into Excel; initial amount of mRNA was calculated and *Cdc25a* mRNA expression was normalized to *Gfp* mRNA. Graphs represent relative *Cdc25a* mRNA expression in relative arbitrary units.

### Immunoblotting

Sperm from ICR (CD-1) males were prepared as in (Fenclová et al, 2022) and were a gift from Jan Nevoral, Charles University, Pilsen, Czech Republic. For immunoblotting, sample amount, collecting time and stage are indicated in the appropriate Figure Legends. Embroys were checked for any remaining cumulus cells and 3x washed in PBS supplied with with 1 mg/ml poly(vinyl alcohol) and only embryos without PBS were freezed at -80°C.

Before immunoblotting, the embryos were lysed in 10-20 µl 1x NuPAGE™ LDS Sample Buffer (Thermo Fisher Scientific, NP0007) supplied with 1x Invitrogen™ NuPAGE™ Sample Reducing Agent (Thermo Fisher Scientific, NP0004) and 1x Halt™ Protease and Phosphatase Inhibitor Cocktail (Thermo Fisher Scientific, 1861281) or Phosphatase Inhibitor Cocktail Set III (Merck, 524627-1ML). The samples were lysed at 100°C for 4 min. Electrophoresis was conducted at 150 V for 4 h on ice in NuPAGE™ MES SDS Running Buffer (20X) (Thermo Fisher Scientific, NP0002) supplied with 1x NuPAGE™ Antioxidant (Thermo Fisher Scientific, NP0005) using NUPAGE 4-12% BT GEL 1.5MM 10W (Thermo Fisher Scientific, NP0335BOX) and Page Ruler PLUS prestained Protein ladder (Thermo Fisher Scientific, 26619). The proteins were blotted using a dry blotting system Trans-Blot Turbo Transfer System (Biorad, 1704150) with Trans-Blot Turbo RTA Mini Nitrocellulose Transfer Kit (Biorad, 1704270). A loading control was performed by total protein staining using Pierce Reversible Protein Stain Kit for Nitrocellulose Membranes (Thermo Fisher Scientific, 24580). For blocking, the membrane was washed in 5% milk in 0.05% Tris-buffer saline-Tween (TBST), pH 7.4 for 1 h. The membrane was then incubated in 1:500 CHK1 (G-4) Antibody (Santa Cruz Biotechnology, sc8408) diluted in 5% milk at 4°C overnight. After 3x washing in TBST for 10 min, the membranes were incubated for 1 h in Peroxidase AffiniPure Donkey Anti-Mouse IgG (Jackson Immuno Research, 715-035-151) diluted 1:5000 in 5% milk in TBST. The membranes were washed 3x in TBST for 10 min again and afterwards incubated in Immobilon ECL Ultra Western HRP Substrate (Merck, WBULS0100) for 2-5 min and scanned with GS-800 densitometer (Bio-Rad).

### Immunofluorescence

Immunofluorescence was performed as described in (Ruth et al, 2021). For quantitative DAPI signal measurements, *Chk1 WT* and *Chk1 mKO* 2-cell stage embryos were initially incubated in 60 µg/ml RNase A Solution (Dynex, #158 922) before DAPI staining in a single experiment without EdU and γH2AX staining. We found the initial RNase treatment was not required and therefore was omitted. Embryos were mounted in ProLong Gold Antifade Mountant with DAPI (Invitrogen, P36941) or laboratory-made Mowiol with 0.25 µg/ml DAPI for DNA staining (Mayer et al, 2016).

For EdU pulse staining, embryos were initially incubated with 100 µM EdU (5-ethynyl-2’-deoxyuridine) (Merck, T511285) for 1 h before washing and fixation. After the permeabilization step, the embryos were incubated in dark at room temperature for 2 h in freshly prepared 10 µM AlexaFluor 488 azide (Invitrogen, A10266) in PBS supplemented with 1 mM copper(II) sulfate (CuSO_4_) (Merck, C1297) and 10 mM sodium-L-ascorbate (Merck, 11140).

### Confocal microscopy

Samples were scanned using a confocal microscope Leica TCS SP5 with with HCX PlApo Lambda Blue 63x 1.4 oil objective. We applied a sequential scan with 400 Hz, 12-bit depth, 512×512 pixels with 120 nm pixel size, zoom 4, line average 3, 1 µm z-sections. TRITC was excited by 561 laser, EdU by 488 laser and DAPI by 405 laser. TRITC and Alexa were scanned using HyD detectors, DAPI was scanned using PMT detector.

### Time-lapse imaging

Bright-field time-lapse imaging was performed on an inverted microscope Leica DMI 6000B (Leica Microsystems, Germany) in a controlled environment (37°C, 5% CO_2_ in air) (del Llano et al, 2022). Oocytes/embryos were placed in a 10 µl drop of culture medium covered with oil (oocytes: Mineral oil, Merck, M5310; embryos: OVOIL™, Vitrolife, 10029). Bright-field images were taken in 30 min intervals. Only embryos with two pronuclei that divided into 2-cell stage were included in the analysis.

Fluorescent long-term time-lapse imaging of embryos was performed on a Viventis LS1 Live light-sheet microscope (Viventis Microscopy Sarl, Switzerland) as described in (Ferencova et al, 2022). Embryos were placed in a 100 µl of culture medium covered with 150 µl oil (OVOIL™, Vitrolife, 10029) in sample holder multi-well (Viventis Microscopy, SHM1_4W). When inhibitors were used embryos were placed in 250 µl culture medium and the dishes were covered with parafilm with small holes to allow gas exchange. Images were taken using a pixel resolution of 750×750 (until the 8-cell stage) or 1024×1024 (until morula/blastocyst stage) with both pixel size 173 nm, 3-μm optical sections and 10-min time intervals. Only embryos with two pronuclei that divided into 2-cell stage were included in the analysis.

Because both 2- to 3- and 3- to 4-stage cell divisions represent second embryonic division, we chose to show 2- to 3-cell stage division. On the other hand, in assessing chromosome segregation errors, we scored both 2- to 3- and 3- to 4-stage cell divisions to represent the entire second embryonic division.

### Image analysis

Image analysis was performed using Fiji software (Schindelin et al, 2012). Cell cycle phases were determined using 60 min EdU pulse staining, temporal information and chromosome condensation state as shown by DAPI staining. EdU-positive embryos were scored as in S phase. EdU-negative embryos were classified being in G2. Mitosis was detected by the state of chromosome condensation from DAPI staining. The number of γH2AX foci were determined by the Findfoci algorithm for Fiji (Herbert et al, 2014). DNA content by DAPI staining was measured as a sum of the pixel values of DAPI signal in the nucleus.

The mCDT1-EYFP, CDC25A-EGFP, and EGFP raw signal was measured on maximal intensity projections in x-axis. Values were normalized to maximum mean value at the peak or maximum values for EGFP. Cell cycle phase lengths from the mCDT1-EYFP sensor were determined as follows: G1 was assessed as the time from nuclear envelope assembly (marked by the nuclear mCDT1-EYFP signal appearance) until the peak value. Initiation of S phase was assessed as the first time-point of a declining signal after the peak value. Completion of S phase was determined as the start of G2. Start of G2 was assessed as the first time-point of the first three successive time-points with rising values. End of G2 was assessed as the time of disappearance of mCDT1-EYFP signal during NEBD. For CDC25A-EGFP, signal disappearance before 2 5h after nuclear assembly indicated NEBD.

For representative videos, the fluorescent images were denoised by Gaussian blur (r=1) and for H2B-mCHERRY was applied an autocontrast macro.

### Statistical analysis

Most data were obtained from at least two independent experiments and the data were pooled. Statistical analysis was performed using NCSS 11 software (NCSS, Kaysville UT, USA). The following test were used: Student’s t-test for normal data distribution with equal variances (unpaired), Aspin-Welch test for normal distribution with unequal variances (unpaired), Mann-Whitney U Test for other continuous distributions, Fisher’s Exact Test for binomial distributions. The results were considered statistically significant with p<0.05 an depicted as *p<0.05, **p<0.01 and ***p<0.001.

## ACKNOWLEDGEMENTS

The authors dedicate this manuscript to the memory of Petr Solc, who directed the lab until his untimely death. D.D also thanks Jan Nevoral for the sperm samples, and Petr Svoboda and Jakub Cervenka for advice. D.D. was funded by grant #20-27742S from the Grant Agency of the Czech Republic. L.K. was supported by Ph.D. grant #1402217 from the Grant Agency of Charles University. V.B. was supported by grant VEGA 2/0072/19 from Grant Agency of Slovakia.

## DATA AVAILABILITY

Bright-field time-lapse images, immunoblots and image analysis of confocal or light-sheet live-cell microscopy are available in the BioStudies database under accession number S-BSST940. Other data are available from the corresponding author upon reasonable request.

## AUTHOR CONTRIBUTIONS

**Lucie Knoblochova:** Conceptualisation; Methodology; Validation; Formal Analysis; Investigation; Data curation; Funding acquisition; Project administration; Visualisation; Writing – original draft; Writing – review & editing. **Tomas Duricek:** Formal Analysis; Investigation; Writing – review & editing. **Michaela Vaskovicova:** Methodology; Software; Investigation; Data curation; Formal Analysis; Resources; Writing – review & editing. **Chrysoula Zorzompokou:** Investigation; Writing – review & editing. **Diana Rayova**: Methodology; Resources; Writing – review & editing. **Ivana Ferencova:** Investigation; Writing review & editing. **Vladimir Baran:** Conceptualisation; Writing – review & editing. **Richard M. Schultz:** Supervision; Writing - review & editing. **Eva R. Hoffmann:** Supervision; Writing review & editing. **David Drutovic:** Conceptualisation; Validation; Project administration; Supervision; Writing – original draft; Writing – review & editing.

## SUPPLEMENTAL FIGURE LEGENDS

**Supplemental Figure 1.**
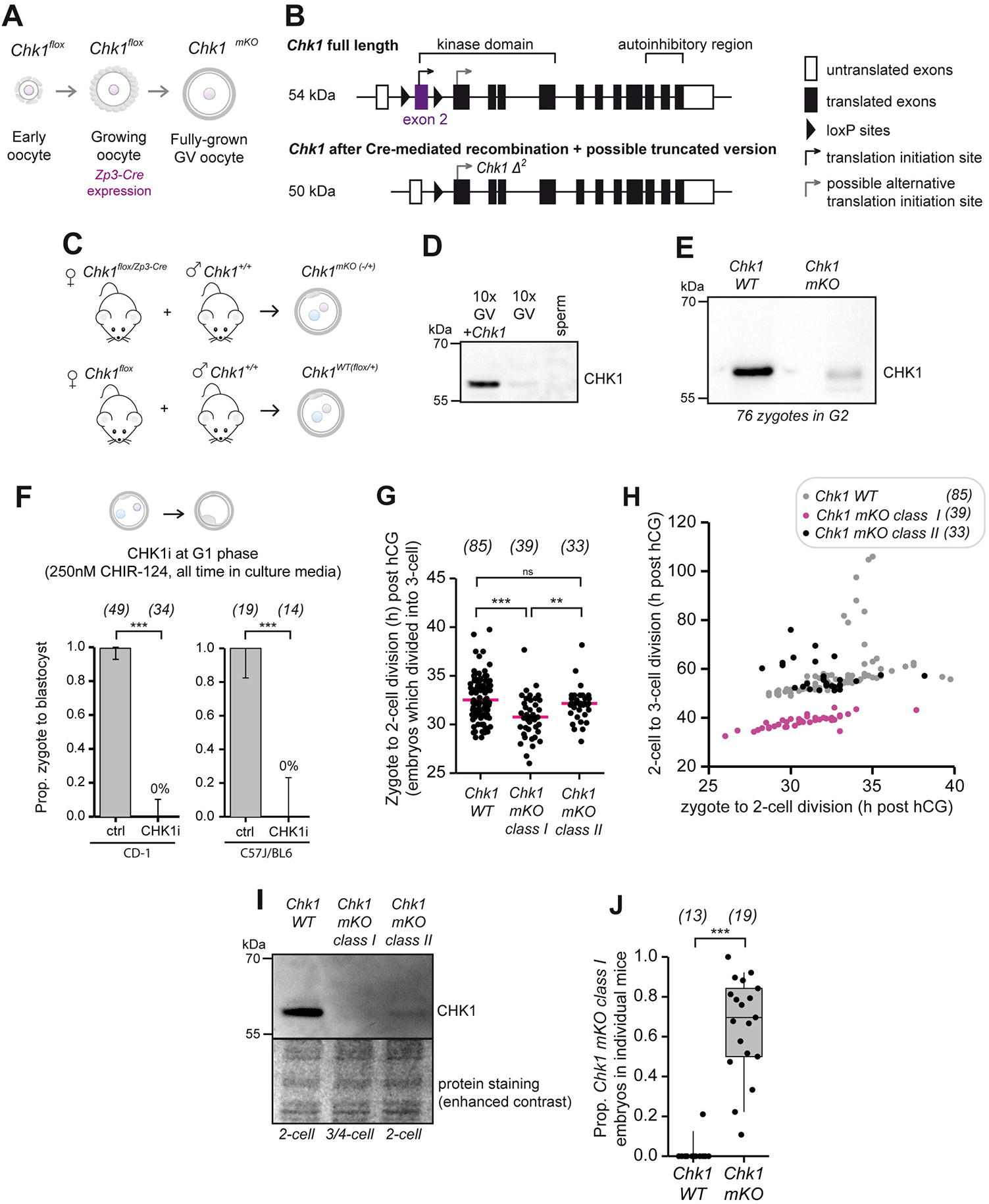
Maternally-expressed CHK1 regulates early embryonic development. **A** Schematic depiction of Cre-recombinase expression specifically in growing oocytes under *zona pellucida 3 (Zp3)* promoter. **B** Schematic depiction of *Chk1* gene with indicated protein domains. Exon 2 of *Chk1*, which contains a translation initiation site and is flanked by LoxP sites, is excised by Cre-recombinase, thereby preventing translation initiation of full-length *Chk1* mRNA. **C** Female mice carrying floxed *Chk1* exon 2 and Cre-recombinase under oocyte-specific *Zp3* promoter are mated with *Chk1* wild-type (WT) male producing maternal *Chk1* knock-out embryos with present paternal allele (*Chk1 mKO*). **D** CHK1 protein level detected by immunoblot from positive controls (10 germinal-vesicle (GV) stage oocytes microinjected with full-length 300 ng/µl of *Chk1* cRNA), 10 GV oocytes (GV WT), and sperm (0.22 × 10^6^). kDa kilodalton. **E** CHK1 protein levels detected by immunoblot in *Chk1 WT* and *Chk1 mKO* zygotes at 26 h post hCG administration. **F** Proportion of zygotes developed to blastocysts after pharmacological CHK1 inhibition in C57J/BL6 or CD-1 females. The numbers in brackets denote the number of analyzed embryos. Error bars denote the 95% confidence interval. ***p< 0.001 (Fisher’s Exact Test, two-sided). **G** Data from bright-field time-lapse imaging (**Fig. 1B**) was used to quantify the division time (h) from zygote to 2-cell in *Chk1 WT, Chk1 mKO class I*, and *Chk1 mKO class II* embryos categorized retrospectively based on their division timing to the 3-cell stage, related to **Fig. 1B-F**. The median is shown. ns non-significant, **p<0.01, and ***p<0.001 (Mann Whitney U Test) **H** Time of the 2-cell to 3-cell division after hCG administration relative to the zygote to 2-cell division, related to **Fig. 1B-F**. *Chk1 mKO class I* embryos show a positive correlation between the division time from zygote to 2-cell and 2-cell to 3-cell (Pearson’s correlation coefficient 0.7932, p<0.0000). *Chk1 WT* embryos show a positive correlation (Pearson’s correlation coefficient 0.4214, p=0.0001) but with some substantial outliers. *Chk1 mKO class II* shows a non-significant negative correlation (Pearson’s correlation coefficient -0.3080, p=0.0812). **I** CHK1 protein levels detected by immunoblot from 2- or 3-4 cell embryos collected 47 h post hCG administration. n*=*79, 79, and 78 embryos for *Chk1 WT, Chk1 mKO class I*, and *Chk1 mKO class II*, respectively. **J** Proportion of *class I* embryos indicated for individual mice. The embryos were scored at 46-47 h post hCG administration. The numbers in brackets denote the number of mice. The overlay with boxplot is shown. ***p<0.001 (Mann Whitney U Test).

**Supplemental Figure 2.**
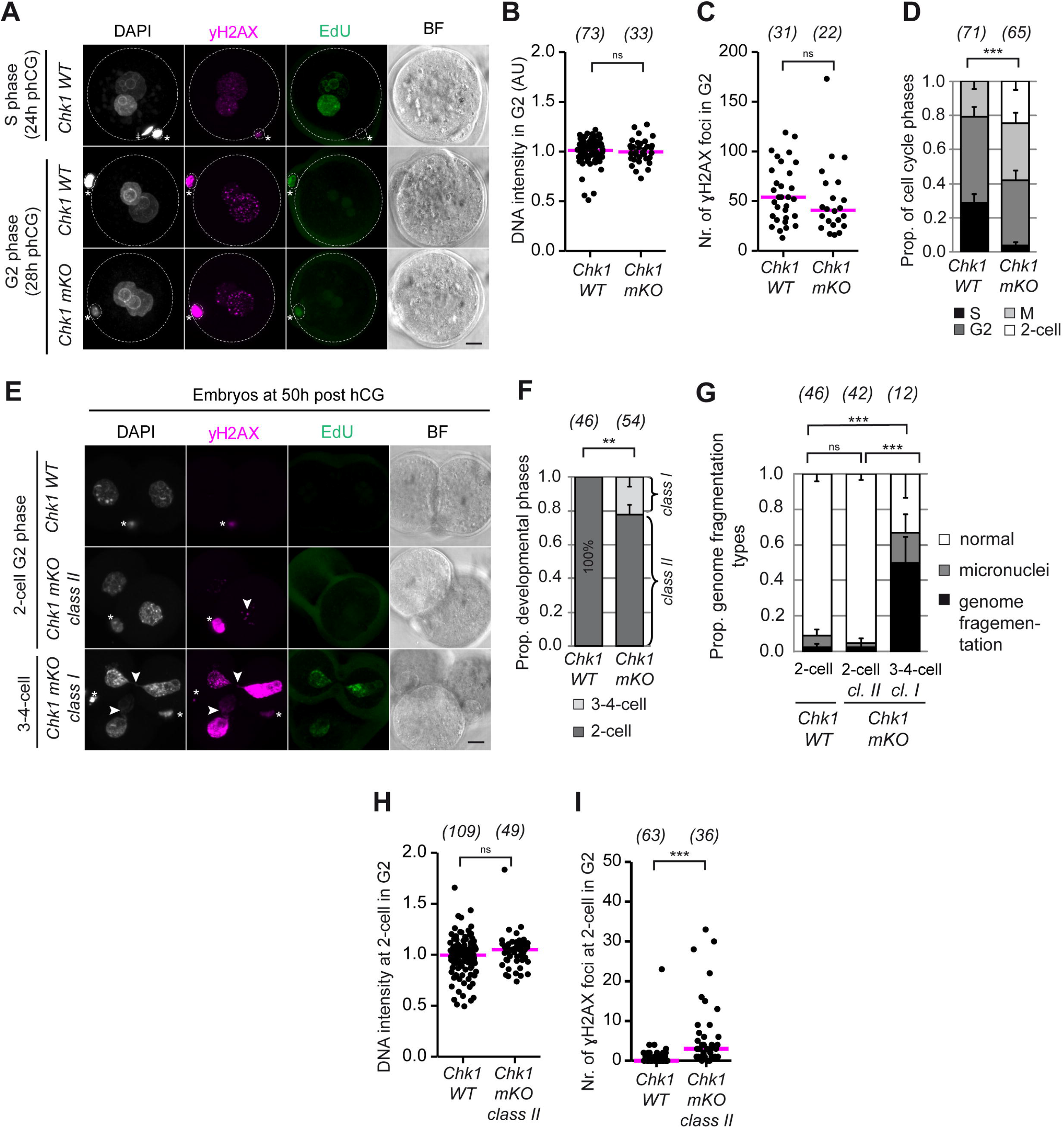
CHK1 kinase regulates cell cycle progression. **A** Representative still images from immunofluorescence analysis of cell cycle distribution in *Chk1 WT* and *Chk1 mKO* embryos combined with γH2AX foci and DNA content analysis. Zygotes in the S phase (24 h post hCG administration) and G2 phase (28 h post hCG administration) were fixed after a 60 min EdU pulse to identify zygotes that completed DNA replication. Embryos with γH2AX signal (magenta), DAPI (chromatin, gray), EdU (green), and BF (bright field) are shown. Asterisk mark polar bodies. Crosses mark the sperm head at the membrane. Scalebar 10 μm. **B, C** Data from immunofluorescence analysis (**A**) was used to quantify DNA content (**B**) and ɣH2AX foci number (**C**) in zygotes in the G2 phase. The median is shown. DNA content was measured by DAPI intensity. The data were normalized to the mean in each group. AU (arbitrary units). The data were pooled from two independent experiments. ns non-significant (Mann-Whitney U Test, two-sided) **D** Data from immunofluorescence analysis (**A**) was used to quantify the proportion of cell cycle phases in embryos. Error bars denote the standard error of the mean. ***p<0.001 (Cochran-Armitage Trend Test, two-sided). **E** Representative still images from immunofluorescence analysis of cell cycle distribution in *Chk1 WT* and *Chk1 mKO* embryos combined with γH2AX foci and DNA content analysis. Embryos at 50 h post hCG administration (G2 phase) were fixed after a 60 min EdU pulse to identify zygotes that completed DNA replication. Embryos with γH2AX signal (magenta), DAPI (chromatin, gray), EdU (green), and BF (bright field) are shown. Asterisk mark polar bodies. Arrowhead points to DNA damage visualized by γH2AX or DAPI. Scalebar 10 μm. **F, G** Data from immunofluorescence analysis (**E**) was used to quantify the proportion of developmental phase (**F**) and genome fragmentation types (**G**) in embryos. Error bars denote the standard error of the mean. The data were pooled from two independent experiments. ns non-significant, **p<0.01, ***p<0.001 (Cochran-Armitage Trend Test, two-sided). **H, I** Data from immunofluorescence analysis (**E**) was used to quantifying DNA content (**H**) and γH2AX foci number (**I**) in embryos at 50 h post hCG administration. Median is shown. DNA content was measured by DAPI intensity. The data were normalized to the mean in each group. AU (arbitrary units). The data were pooled from two independent experiments. ns non-significant, ***p<0.001 (Mann-Whitney U Test, two-sided).

**Supplemental Figure 3.**
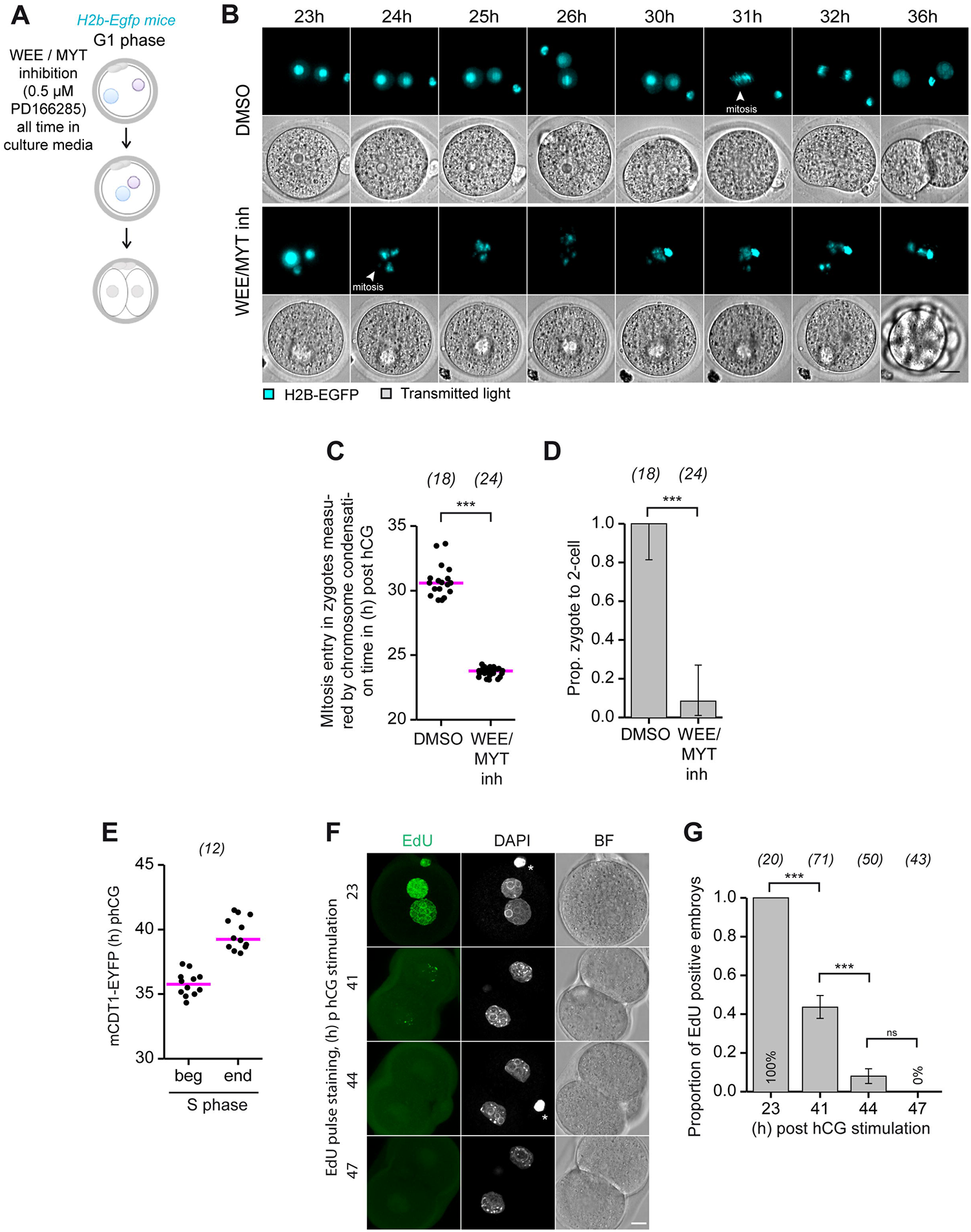
Characterisation of replication checkpoint regulation and duration of S-phase. **A** Schematic of the experimental procedure. **B** Representative still images from time-lapse light-sheet imaging of embryos from *H2b-Egfp* transgenic mice from 23 to 36 h post hCG administration after WEE/MYT kinase inhibition (as shown in **A**) or DMSO treatment (21 h post hCG administration). Embryos expressing H2B-EGFP (cyan, chromosomes) are shown. Transmitted light (grays). Arrows indicate mitotic entry. Scalebar 20 μm. Supplemental Movie 1 related to Supplemental Fig. S3B: ctrl WEE/MYT inhibition **C** Data from time-lapse light-sheet imaging (**B**) was used to quantify chromosome condensation time (h) from zygote to 2-cell stage embryos. The data were pooled from two independent experiments. Median is shown. ***p<0.001 (Mann-Whitney U Test, two-sided). **D** Data from time-lapse light-sheet imaging (**B**) was used to quantify the proportion of zygote to 2-cell development. The data were pooled from two independent experiments. Error bars denote the 95% confidence interval. ***p<0.001 (Fisher’s Exact Test, two-sided). **E** Beginning and ending of S phase measured by mCDT1-EYFP sensor fluorescence signal in wild-type embryos using time-lapse light-sheet imaging. The magenta bar represents the median of the S phase beginning and ending. **F** Representative images from analysis of S phase estimated by EdU pulse staining in wild-type embryos at 23, 41, 44, and 47 h post hCG administration. The embryos were fixed after a 60 min EdU pulse (green), and DNA was visualized by DAPI staining (grays, chromatin). BF (bright field). Asterisk mark polar bodies. Scalebar 10 μm. **G** Data from immunofluorescence analysis (**F**) was used to quantify the proportion of EdU-positive embryos. Error bars denote the standard error of the mean. The data are from one experiment. ns non-significant, ***p<0.001 (Fisher’s Exact Test).

**Supplemental Figure 4.**
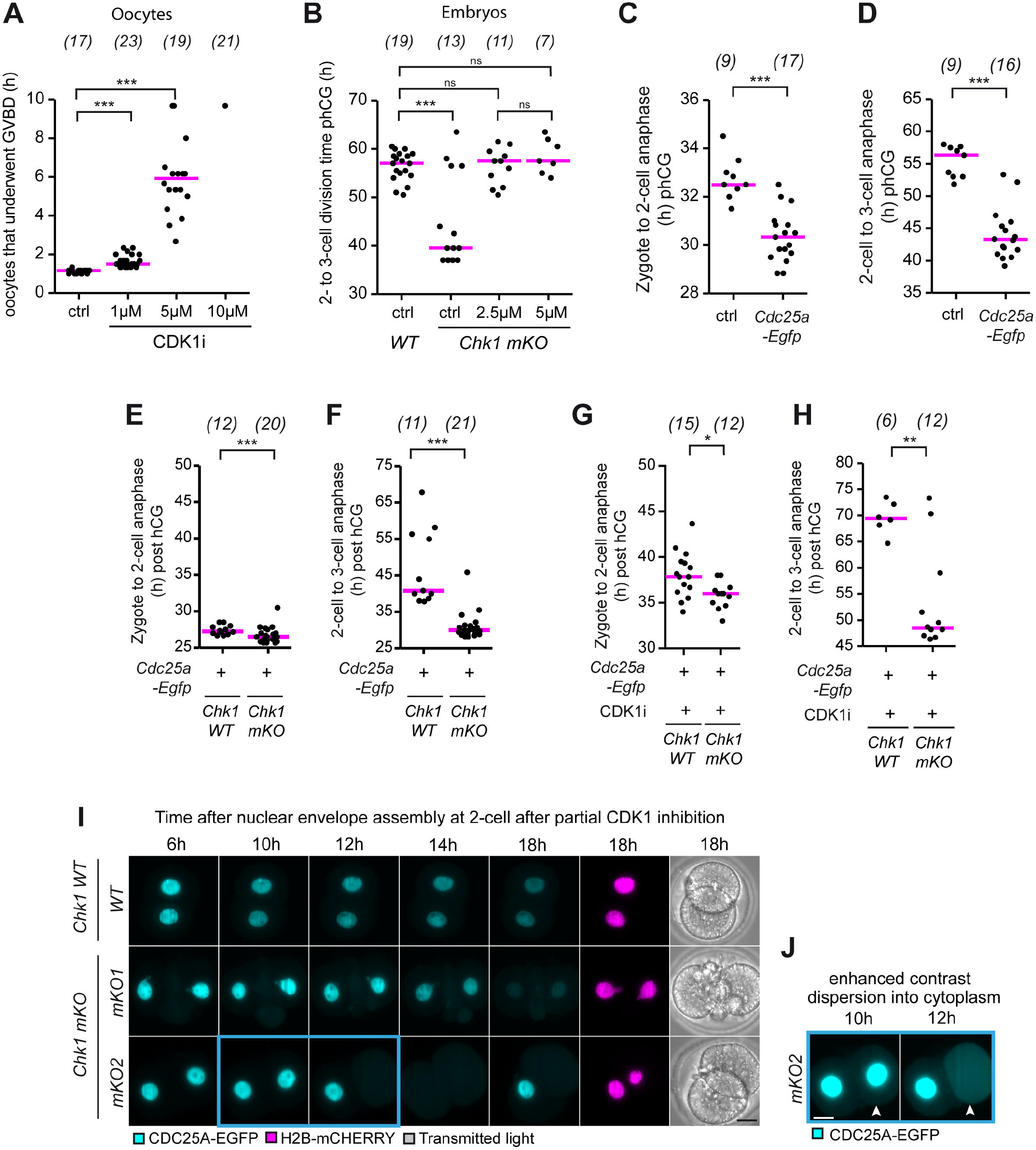
CHK1 kinase regulates the long G2 phase in 2-cell embryos by restraining CDK1 activity via CDC25A degradation. **A** Dot-plot graph of GVBD time (h) analyzed from time-lapse wide-field imaging of oocytes after transfer to milrinone-free medium and treated with CDK1 kinase inhibitor (1, 5 and 10 μM RO3306). The numbers in parentheses denote the number of analyzed oocytes per group. In the 10 μM RO3306 group, only 1 oocyte from 21 resume meiosis. The data are from one experiment. ***p<0.001 (Mann-Whitney U Test). **B** Dot-plot graph of division time (h) from 2-cell to 3-cell stage analyzed from time-lapse wide-field imaging of wild-type (WT) and *Chk1 mKO* embryos. The *Chk1 mKO* embryos were treated with 2.5 or 5 µM RO3306. Strain background: CD-1 with small part C57J/BL6 and C57J/BL6 with small part CD-1. ns non-significant, ***p<0.001 (Kolmogorov-Smirnov Test) **C, D** Data from time-lapse light-sheet imaging was used to quantify the initiation of anaphase (h) from zygotes to 2-cell stage embryos (**C**) and from 2-cell to 3-cell embryos (**D**) in WT embryos microinjected with *Cdc25a-Egfp* cRNAs. The data were pooled from two independent experiments. Median is shown. ***p<0.001 (Mann-Whitney U test, two-sided). **E, F** Data from time-lapse light-sheet imaging **(Fig. 3B)** was used to quantify initiation of anaphase (h) from zygote to 2-cell (**E**) and from 2-cell to 3-cell (**F**) in *Cdc25a-*microinjected *Chk1 WT*, and *Cdc25a-*microinjected *Chk1 mKO*. Median is shown. *p<0.05, ***p<0.001 (Mann-Whitney U test, two-sided). **G, H** Data from time-lapse light-sheet imaging (**Fig. 3E**) was used to quantify initiation of first anaphase time (h) (**G**) and time of entry into the second mitosis (**H**) in *Chk1 WT* and *Chk1 mKO*, embryos microinjected with *Cdc25a-Egfp* cRNA and treated with CDK1 inhibitor. Due to experimental procedures, not all embryos underwent cytokinesis after the first mitosis. Median is shown. *p<0.05, **p<0.01 (Mann-Whitney U test, two-sided). **I** Representative still images from time-lapse light-sheet imaging (**Fig. 3E**) of *Chk1 WT* and *Chk1 mKO* embryos treated with CDK1 inhibitor. Embryos were microinjected, as shown in **Fig. 3E**. Embryos expressing H2B-mCHERRY (magenta, chromosomes) and CDC25A-EGFP (cyan) are shown. Scalebar 20 μm. **J** Enhanced contrast of still images of mKO2 at 10 h and 12 h. Arrowheads mark CDC25A-EGFP signal in cytoplasm. Note that cytoplasmic signals increase from 10 to 12 h indicating that there is CDC25A-EGFP dispersed from the nucleus into cytoplasm (not degradation). Supplemental Movie 3 related to Supplemental Fig. S4I, J and Fig. 3F, G *Chk1 WT* + CDC25A-EGFP + CDK1i *Chk1 mKO (mKO1)* + CDC25A-EGFP + CDK1i *Chk1 mKO (mKO2)* + CDC25A-EGFP + CDK1i

**Supplemental Figure 5.**
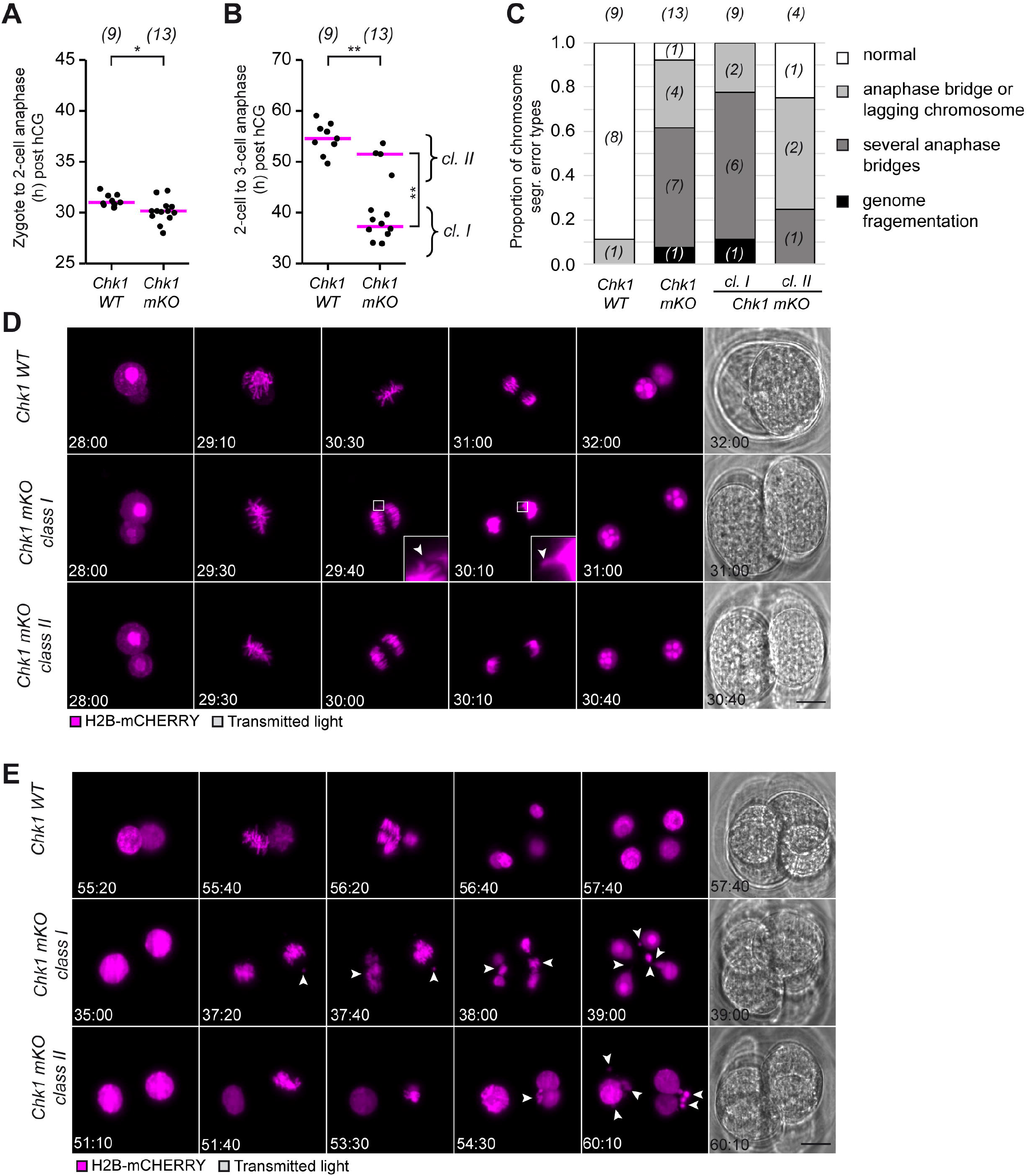
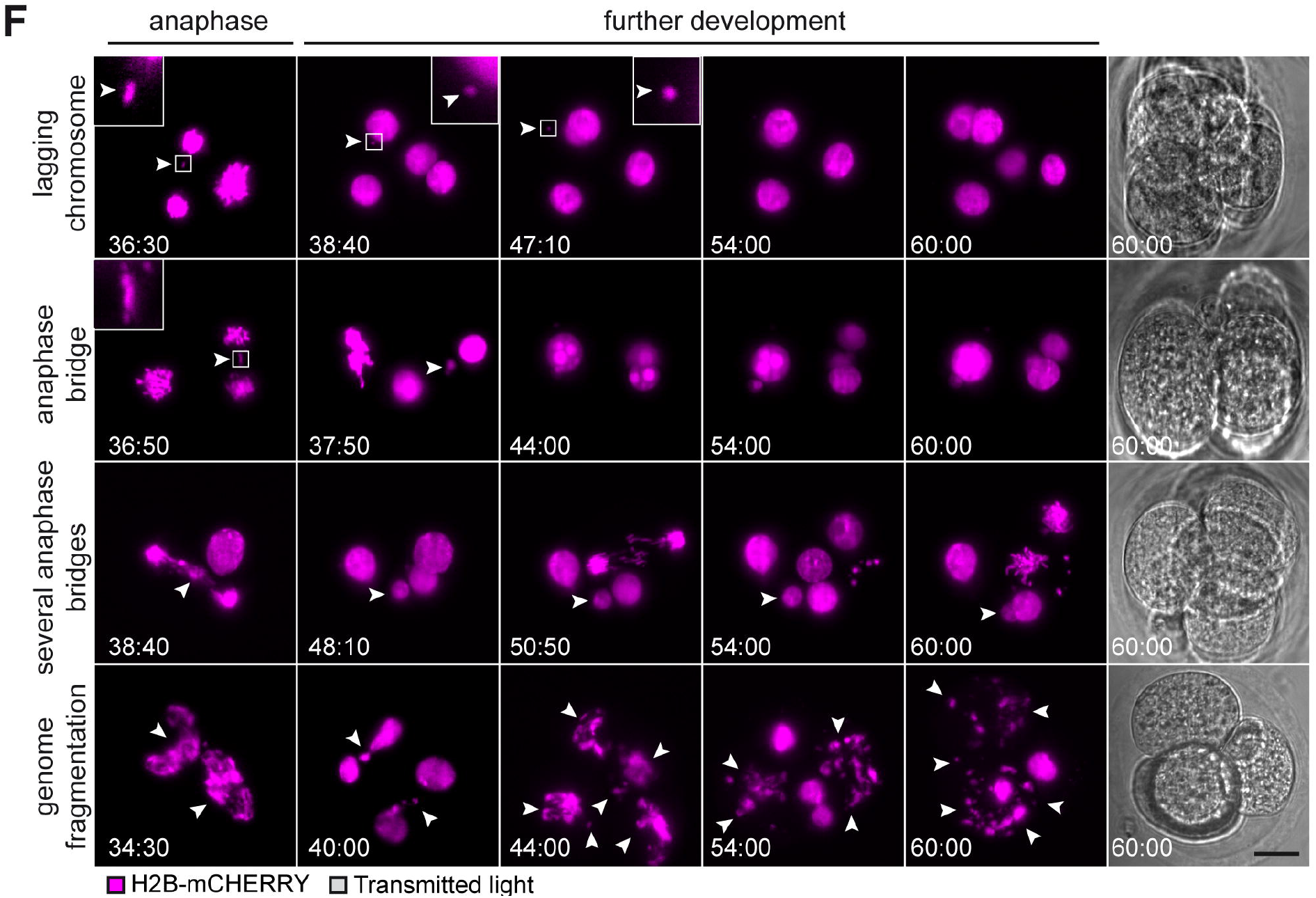
CHK1 is essential for DNA integrity protection in the second embryonic division. **A, B** Data from time-lapse light-sheet imaging (**Fig. 5B**) was used to quantify initiation of anaphase (h) from zygote to 2-cell (**A**) and time of division (h) from 2-cell to 3-cell (**B**) in *Chk1 WT* and *Chk1 mKO* embryos. A subgroup of *Chk1 mKO* embryos (C*hk1 mKO class I*) shows substantially accelerated second division. The numbers in brackets denote the number of analyzed embryos per genotype. The median is shown. The data were pooled from 2 independent experiments. *p<0.05 (Equal-Variance T-Test, two-sided). *p<0.05 relative to class I (Kolmogorov-Smirnov Test, two-sided), *p<0.05 relative *Chk1 WT* (Mann-Whitney U Test). **C** Data from time-lapse light-sheet imaging (**Fig. 5B**) was used to quantify the proportion of different types of chromosome segregation errors in embryos. **D, E, F** Representative still images from time-lapse light-sheet imaging (**Fig. 5B**) of the first embryonic division (**D**), second embryonic division (**E**), and different chromosome segregation error types (**F**) in wild-type (*Chk1 WT*), *Chk1 mKO class I* and *Chk1 mKO class II* embryos. Embryos were microinjected, as shown in **Fig. 5A**. Embryos expressing H2B-mCHERRY (magenta, chromosomes) are shown. Scalebar 20 μm. Supplemental Movie 6 related to Supplemental Fig. S5D, E, F: lagging chromosome anaphase bridge several anaphase bridges genome fragmentation

**Suppl. Figure 6.**
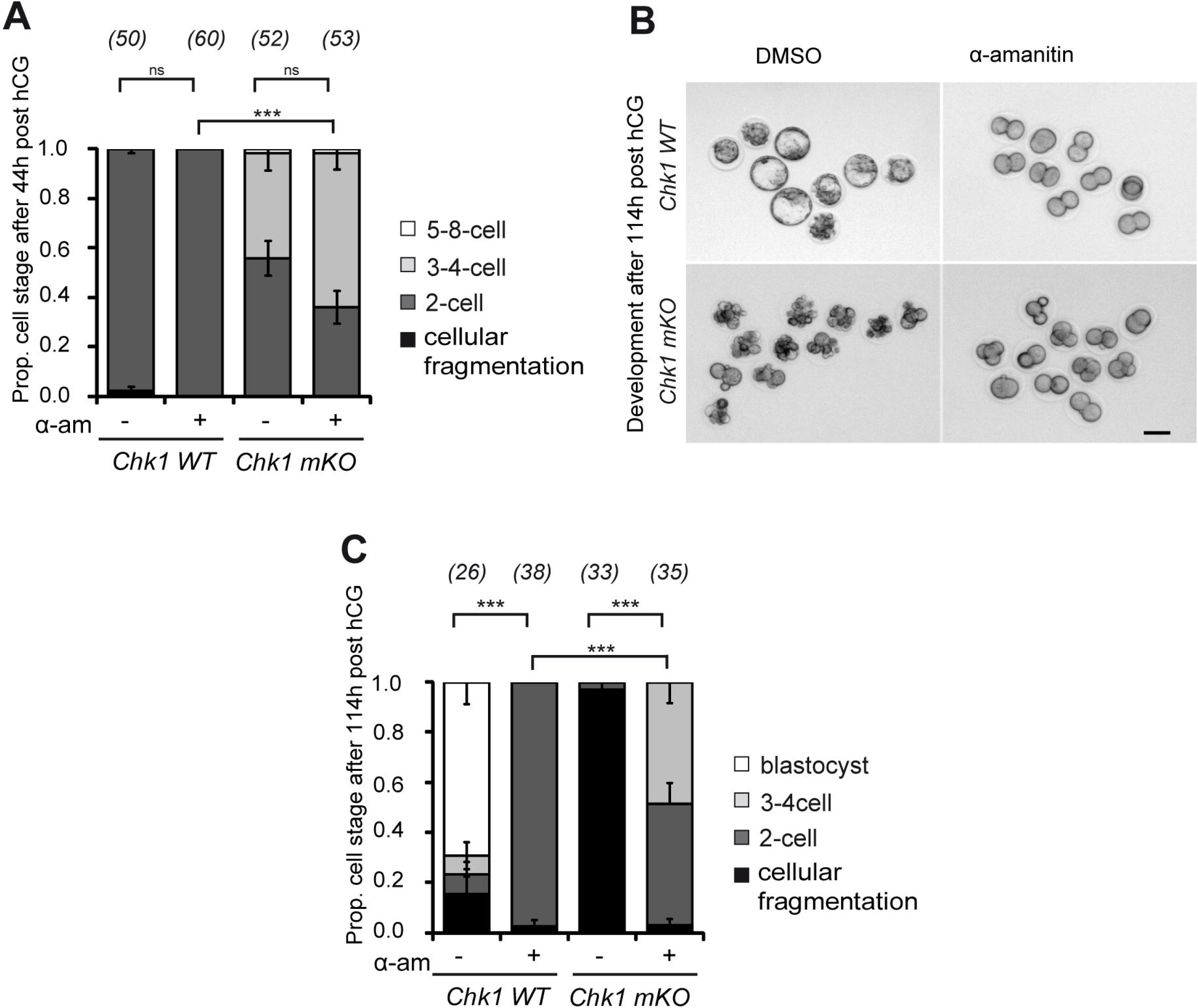
CHK1 is necessary for the 2-cell stage transcription block. **A** Developmental analysis of *Chk1 WT* and *Chk1 mKO* embryos treated with α-amanitin (24 µg/ml) or DMSO. The embryos were scored at 44 h post hCG administration. The numbers in parentheses denote the number of analyzed embryos per group. The data were pooled from two independent experiments. Strain background: CD-1 with small part C57J/BL6 and C57J/BL6 with small part CD-1. Error bars denote the standard error of the mean. ns non-significant, ***p<0.001 (Cochran-Armitage Trend Test, two-sided. **B** Representative images from developmental analysis (**C**) of developmental stages at 114 h post hCG administration. Scalebar 60 μm. **C** Developmental analysis of *Chk1 WT* and *Chk1 mKO* embryos treated with α-amanitin (24 µg/ml) or DMSO. The embryos were scored at 114 h post hCG administration. The numbers in parentheses denote the number of analyzed embryos per group. The data were pooled from two independent experiments. Error bars denote the standard error of the mean. ***p<0.001 (Cochran-Armitage Trend Test, two-sided.

